# Non-catalytic role for MLL2 in controlling chromatin organisation and mobility during priming of pluripotent cells for differentiation

**DOI:** 10.1101/2025.02.21.639010

**Authors:** Maike Steindel, Oliver Davis, Katrin Neumann, Gökçe Agsu, Lingjun Mao, Andrea Kranz, Liviu Pirvan, Dwaipayan Adhya, Jorg Morf, Ziwei Zhang, Shuyue Yang, Jun Fu, Melania Barile, Annabelle Wurmser, Stanley E. Strawbridge, Nvard Chalabyan, Pradeepa Madapura, Brian Huntly, Berthold Göttgens, David Holcman, Shamith A. Samarajiwa, David Klenerman, Konstantinos Anastassiadis, A. Francis Stewart, Srinjan Basu

## Abstract

How the chromatin regulator MLL2 (KMT2B) influences cell differentiation remains poorly understood. MLL2 is the main histone 3 lysine 4 (H3K4) trimethyltransferase acting at bivalent promoters in embryonic stem cells (ESCs) and is required for ESCs to differentiate into neuroectoderm. We show here that this requirement occurs during exit from naïve pluripotency, days before neuroectoderm differentiation is impaired. Although MLL2 knockout has only a subtle effect on transcription during exit, reducing the expression of a few important neuroectodermal transcription factors, it substantially remodels chromatin architecture, disrupting 3D chromatin loops associated with bivalent promoters. The enzymatic activity of MLL2 is not needed for stabilising these loops or for neuroectoderm differentiation. This non-catalytic function of MLL2 in stabilising 3D chromatin loops has implications for lineage specification. Because MLL2 shares features with all four MLLs, chromatin tethering, rather than H3K4 methylation, may represent the primary function of MLL proteins during lineage commitment.

## Introduction

Multicellularity occurs because stem cells can utilize information encoded in the genome in alternative ways to differentiate into distinct cell types. These capacities involve the hierarchical regulation of gene expression by transcription factors coupled with epigenetic hierarchies, including alterations in 3D chromatin organisation (*1–5*). Together these mechanisms elicit the multiple utilizations of the genome required for multicellularity. Unravelling these molecular mechanisms is key to understanding development, disease, and the therapeutic potential of stem cells for regenerative medicine.

The differentiation of mammalian pluripotent cells, including embryonic stem cells (ESCs), both *in vitro* and *in vivo*, has revealed a multi-step process in which naïve ESCs generate primed ESCs before differentiation toward cell lineages such as neuroectoderm. *In vivo*, naïve pluripotent cells that emerge in the epiblast of pre-implantation embryos (between embryonic day E3.75 and E4.75 in the mouse (*6*)) have the potential to differentiate into all cell lineages of the mammalian body (*7*). Pluripotent cells transition from a naïve to a primed pluripotent state as the embryo implants (between E4.5 and E6.5) (*8, 9*). *In vitro*, ESCs derived from the inner cell mass of pre-implantation blastocysts are equivalent to naïve pluripotent cells when cultured under defined conditions called 2iLIF (*10*). Removal of 2iLIF promotes exit from naïve pluripotency (*11–13*). Subsequent addition of FGF2 and activin promotes priming to post-implantation epiblast-like cells (EpiLCs) (*14–16*). EpiLCs can then be differentiated towards specific lineages such as neuroectoderm following specific inductive cues (*17*).

During the initial priming phase as ESCs exit from self-renewal, the enhancers and/or promoters of lineage-specifying genes become accessible, acquire specific chromatin marks (DNA hypomethylation, H3K4me1/3, H3K27ac/me3) (*18–21*) and alter their 3D genome architecture (*22, 23*). Depleting proteins responsible for these changes perturbs differentiation, supporting the proposition that early changes in the chromatin landscape prime ESCs towards specific lineages (*18–21, 23*).

Amongst chromatin regulators, MLL1-4 (also termed KMT2A-D) play central roles in embryonic development, differentiation and stem cell maintenance in a range of mammalian cell lineages (*24*). In particular, MLL1 is required for three types of adult stem cell in mammals: hematopoietic stem cells, intestinal stem cells and satellite cells in skeletal muscle (*25–27*). Notably, the MLLs are histone 3 lysine 4 methyltransferases (H3K4 MTs). Despite the relevance of H3K4 methylation to epigenetic regulation (*28*), the methyltransferase activities of the MLLs are not always required for the critical functions played by these proteins (*29–31*). In all four MLLs, the C-terminal SET methyltransferase domain is only a small part of these large nuclear proteins (MLL2 is ∼300 kDa). Notably, MLLs also contain arrays of chromatin binding zinc fingers (PHD and CXXC), intra- (FYRN, FYRC) and intermolecular protein-protein interaction domains including a CBP/p300 transactivation region, interspersed amongst extensive intrinsically disordered regions (IDRs) (*24*). Indeed, in higher eukaryotes the MLLs and orthologues are all large proteins, even though most of them are IDRs that are highly variable. The MLLs clearly convey functions beyond their role as methyltransferases that warrant further investigation.

Here, we address how MLL2 and its catalytic activity influence neuroectoderm differentiation. Although dispensable for self-renewal of naïve ESCs, MLL2 is required for the coordination and timing of differentiation (*32–34*). MLL2 is the H3K4 MT primarily responsible for H3K4me3 deposition at bivalent promoters in ESCs. Accordingly, MLL2 is bound at most bivalent promoters. However, MLL2 also binds at most active promoters and is not required for the vast majority of inducible gene expression (*35*). Notably, MLL2 opposes silencing by Polycomb-Group action and DNA methylation (*34, 36, 37*) and also influences DNA accessibility and 3D genome organisation (*30*). But how do MLL2 and its catalytic activity influence neuroectoderm differentiation? To address this, we combined single-cell transcriptomics, chromosome conformation capture (Micro-C) and single-molecule localisation microscopy (SMLM) to establish that MLL2 functions as naïve ESCs transition into primed ESCs. MLL2 acts by stabilising enhancer-promoter loops, particularly at bivalent genes, independent of H3K4 trimethylation. This stabilisation appears to be MLL2’s primary function in early neuroectoderm lineage determination.

## Results

### The requirement for MLL2 in neuroectoderm differentiation is exerted during ESC exit from naïve pluripotency

A requirement for MLL2 in neuroectoderm differentiation has previously been identified (*33*). To determine when MLL2 is required, we used ligand-regulated conditional mutagenesis (*38*) and *Mll2* conditional knock-out (cKO) ESCs (Denissov et al., 2014) to delete *Mll2* at different timepoints during priming and neuroectoderm differentiation (**Figure 1A-E and S1**). We monitored neuroectoderm differentiation using a range of methods: phase contrast microscopy of neurospheres and neural stem cells (**Figure 1C-D and S1B**) and immunofluorescence of several neural lineage marker genes (NESTIN, PAX6, ZO-1) in neural rosettes (**Figure 1E and S1C-D**). This revealed that the requirement for MLL2 is exerted in the earliest possible window during ESC exit, which is about three days and >6 cell cycles before the emergence of the differentiation defect (**Figure 1D**). To ensure specificity, ectopic expression of GFP-tagged MLL2 from a BAC (bacterial artificial chromosome) transgene rescued neuroectoderm differentiation of *Mll2* cKO cells (**Figure 1B-E and S1B-C**). Because tamoxifen-induced cKO led to loss of MLL2 protein in ESCs within 48 hours (**Figure S1A**), MLL2 must be acting within the first 4 cell cycles during the ESC exit from self-renewal to influence subsequent neuroectoderm differentiation. Notably, cells deleted for *Mll2* at the EpiLC stage (**Figure 1F**) were unimpaired in neuroectoderm differentiation, again indicating that MLL2 is no longer required after ESC exit and exerts its function either during ESC self-renewal, exit, and/or priming several days before the major defects in neuroectodermal differentiation are observed.

**Figure 1.**
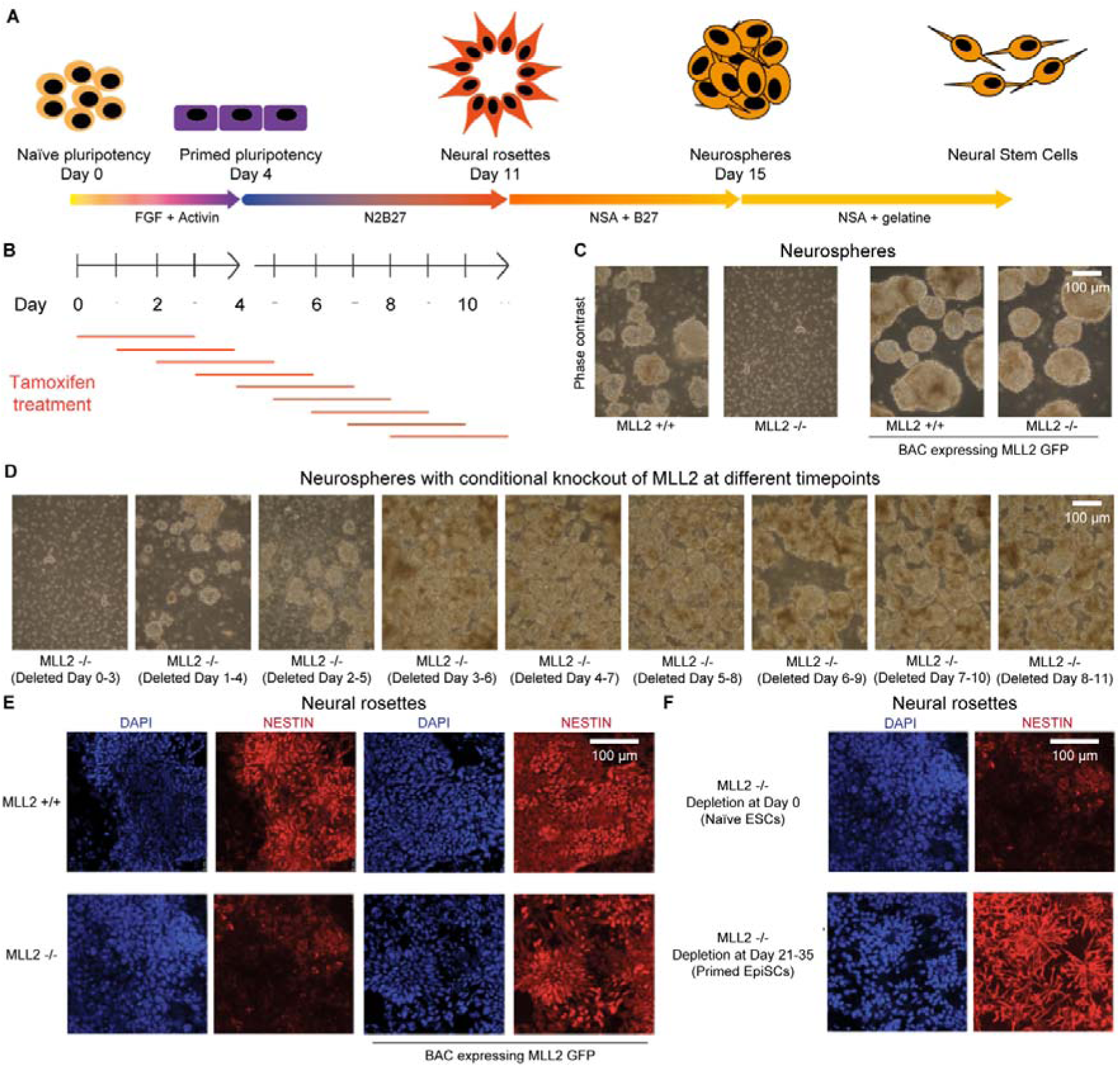
MLL2, which is required for neuroectoderm differentiation, exerts its effect during a priming window. **(A)** Schematic showing protocol for priming of naïve embryonic stem cells (ESCs) and subsequent neuroectoderm differentiation. **(B)** Time course strategy for tamoxifen induced conditional knockout (cKO) of *Mll2*. **(C)** Phase contrast images at the neurosphere stage of control (MLL2 +/+) and *Mll2* cKO (MLL2−/−) cells show that *Mll2* cKO cells fail to generate neural stem cells or neurospheres. Expression of exogenous GFP-tagged MLL2 from an integrated bacterial artificial chromosome (BAC) rescues failure to form neural stem cells and neurospheres in cells lacking endogenous *Mll2* (n=3). **(D)** Phase contrast images at the neurosphere stage from cells subjected to tamoxifen-induced *Mll2* cKO at different days during neuroectoderm differentiation as indicated above and in **Figure S1D**. **(E)** Fluorescence images of DAPI stain (blue) and NESTIN expression (red) at the neural rosette stage of control (MLL2 +/+) and *Mll2* cKO (MLL2−/−) cells show that *Mll2* cKO cells fail to generate neural rosettes while ectopic expression of exogenous MLL2 rescues this failure. **(F)** MLL2 is required prior to priming for successful ES cell differentiation to neural rosettes. Fluorescence images of DAPI stain (blue) and NESTIN expression (red) at the neural rosette stage of control (MLL2 +/+), *Mll2* cKO cells where tamoxifen-induced MLL2 depletion is carried out at Day 0 (MLL2−/− naïve ESCs) or after 21-35 days of FGF-Activin addition (MLL2−/− primed EpiSCs).

### Both wild-type and MLL2 knockout ESCs transition from naïve to primed pluripotency via an intermediate stage with features of formative pluripotency

The tamoxifen cKO time course indicated that, for neuroectoderm differentiation, MLL2 is required before the ESC transition from naïve to primed pluripotency is completed. To gather further evidence about this transition, we used single-cell RNA-seq (scRNA-seq) to compare transcription in wild-type (control) and *Mll2* cKO cells every 24 hours for 3 days during the priming transition (**Figure 2A**). scRNA-seq was employed because priming occurs through intermediate cell states with some asynchronicity (*39–42*). In total, 8737 single-cell transcriptomes (4597 *Mll2+/+,* 4140 *Mll2−/−*) passed stringent quality control measures (**Figure S2A**). As replicates varied in sequencing depth/cell, with more unique transcripts and genes detected per cell from replicate 1 than from replicate 2 (**Figure S2B-C**), high-confidence conclusions were drawn only when both replicates showed similar trends.

**Figure 2.**
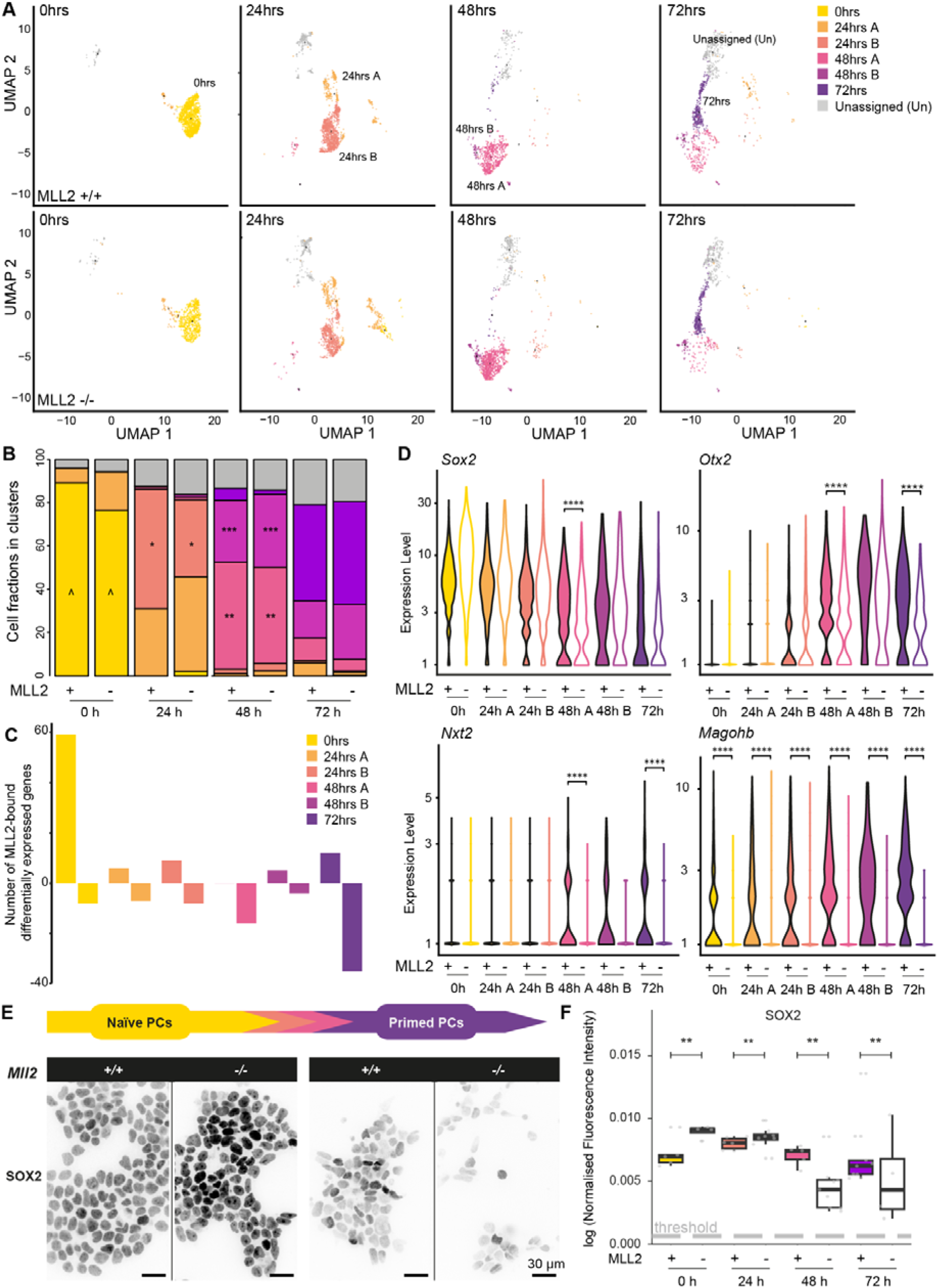
MLL2 knockout has minor effect on cell types generated during priming but reduces expression level of key neuroectoderm-specifying genes in primed pluripotent cells. **(A)** Uniform manifold approximation and projection (UMAP) plot showing wild-type (WT) and *Mll2* cKO (KO) cells during transition from naïve to primed pluripotency with cells collected from each time point and sample shown on separate plots. Cells are coloured differently if they have been assigned to different clusters (see **Methods**). This shows that WT and KO cells both contribute to all assigned clusters with one/two clusters predominating each timepoint. Number of cells: 1215/638 (0 hr +/− MLL2), 428/532 (24 hr A +/− MLL2), 998/671 (24 hr B +/− MLL2), 759/914 (48 hr A +/− MLL2), 133/223 (48 hr B +/− MLL2), 348/343 (72 hr +/− MLL2) and 716/819 (Unassigned +/− MLL2). **(B)** Fraction of cells in WT and KO cells show initial differences in pluripotent cell types that resolve into similar primed pluripotent cell types (^p = 2.48*10^−6^, *p = 0.004, **p = 0.043, ***p = 0.011, Fisher’s exact test). Number of cells: 1308/700 (0 hr +/− MLL2), 1486/1335 (24 hr +/− MLL2), 817/1271 (48 hr +/− MLL2), 986/834 (72 hr +/− MLL2). **(C)** Bar plot of the number of up-/down-regulated differentially expressed genes (DEGs) when comparing WT and KO cells for each cell cluster identified in Figure 2 (p < 0.001, log_2_FC > 0.5 or log_2_FC < −0.5). Colours depict clusters identified in Figure 2. Genes are filtered to ensure they have MLL2 bound within 1 kb of the gene transcription start site. (D) *Mll2* cKO reduces *Sox2* and *Otx2* transcription during priming. Violin plots generated from WT and KO cells show the transcript level in each cluster for the DEGs *Sox2*, *Otx2*, *Nxt2* and *Magohb*. Number of cells: 1215/638 (0 hr +/− MLL2), 428/532 (24 hr A +/− MLL2), 998/671 (24 hr B +/− MLL2), 759/914 (48 hr A +/− MLL2), 133/223 (48 hr B +/− MLL2), 348/343 (72 hr +/− MLL2). [Wilcoxon rank sum tests with Bonferroni correction for *Sox2*, p = 10^−4^ (48 hr A +/− MLL2); for *Otx2*, p = 10^−11^ (48 hr A +/− MLL2), p = 10^−9^ (72 hr +/− MLL2); for *Nxt2*, p = 10^−28^ (48 hr A +/− MLL2), p = 10^−8^ (72 hr +/− MLL2); for *Magohb*, p = 10^−52^ (0 hr +/− MLL2), p = 10^−64^ (24 hr A +/− MLL2), p = 10^−37^ (24 hr B +/− MLL2), p = 10^−146^ (48 hr A +/− MLL2), p = 10^−26^ (48 hr B +/− MLL2), p = 10^−68^ (72 hr +/− MLL2)]. (E) *Mll2* cKO reduces SOX2 protein levels in primed pluripotent cells. Representative immunofluorescence images of SOX2 at 0 and 72 hours of priming in WT and KO cells. **(F)** Boxplot of SOX2 expression at 0, 24, 48 and 72 hours of priming coloured by timepoint. Mean fluorescence per cell is shown as grey dots for each field of view and background threshold as a grey dotted line. Number of images (∼10-100 cells/image): 6/4 (0 hr +/− MLL2), 6/10 (24 hr +/− MLL2), 5/10 (48 hr +/− MLL2), 11/4 (72 hr +/− MLL2). [unpaired two-sided t-test, p = 0.007 (0 hr +/− MLL2), p = 0.008 (24 hr +/− MLL2), p = 0.007 (48 hr +/− MLL2), p = 0.007 (72 hr +/− MLL2).]

Using these datasets and by conducting cluster stability tests, we identified 7 major cell clusters (**Figure S2D-G**) (*43*). Both wild-type and *Mll2* cKO cells contributed to all 7 clusters (**Figure 2A-B**). Although only one major cell population was detected for naïve ESCs at 0 hours (“0hrs”), there was an increase in co-existing cell clusters during priming with two dominant clusters at 24 and 48 hours (“24hrs A/B” and “48hrs A/B”), and one major cluster at 72 hours (“72hrs”) (**Figure 2A-B**). One unassigned cluster, found at all time points, was excluded from further analysis due to its high levels of apoptotic genes (“unassigned”) (**Figure 2B and S3A**). Clustering was not driven by the cell cycle: although subtle cell cycle differences were observed between wild-type and *Mll2* cKO cells (**Figure S3B**) regressing out cell cycle factors did not affect clustering (**data not shown**).

To functionally assign the clusters, we calculated the Spearman correlation of all genes assigned to the second highest expression quartile (**Figure S3C**). Clusters at 0hrs and 48-72hrs were consistent with prior knowledge of the priming transition, expressing naïve pluripotency factors *Nanog* and *Klf4* at 0hrs and primed genes *Otx2* and *Pou3f1* at 48-72hrs (**Figure S3D-F**) (*11, 12, 14, 15, 44, 45*). Due to these similarities, we concluded that the 0hr cluster represented “naïve” ESCs and the 48-72hrs clusters represented “primed” ESCs. The two 24hrs clusters expressed a unique set of genes, showed strong correlation coefficients (>0.8) and correlated similarly to preceding and later time points (**Figure S3C-D**), suggesting that these 2 cell states were different to both naïve and primed ESCs. Gene expression profiles of the two 24hrs clusters were characterised by increased expression of a unique set of transcription factors (e.g. *Zic3, Tcf15*, *Nr0b1, Dppa2, Dppa4*) and chromatin regulators (e.g. *Rbpj, Jarid2, Dnmt3l*) (**Figure S3D-F**). *Zic3, Rbpj* and *Jarid2* play a role in the exit from naïve pluripotency (*5, 46, 47*), suggesting that cells must transition through this intermediate phase to form primed ESCs (48-72hr). Indeed, UMAP plots generated for RNA trajectory analysis (after excluding the “unassigned” cluster) confirmed that this was the case (**Figure S3G**).

To validate our clusters, we compared our dataset to published *in vivo* and *in vitro* datasets. *In vivo* datasets (*48*) revealed that our transition mimics the *in vivo* pre-to-post-implantation embryo transition from the E3.5 to E6.5 epiblast (**Figure S3H**). Next, we assessed whether our 24hrs clusters were linked to the hypothesis of a transient state between naïve and primed pluripotency called formative pluripotency, a state with higher multi-lineage potential than both naïve and primed ESCs (*39*). Indeed, expression of *Dppa2/*4 and *Tcf15* during this transition have previously been shown to correlate with higher multi-lineage potential (*19, 49*), lending support to this hypothesis. We therefore compared our data to cells recently captured *in vitro* that have features of formative pluripotency (*40*) and found that they were a mixture of our 24hrs and 48hrs clusters (**Figure S3I**). These comparisons suggested that our datasets were relevant to *in vivo* cell types and that our 24hr and 48hr clusters had features of formative pluripotency.

### MLL2 knockout has minor transcriptional impact during the priming of pluripotent cells, reducing the expression of a few key neuroectoderm genes

Cell cluster identities at 72hrs remained largely unchanged in *Mll2* cKO cells compared to control cells (p > 0.1) (**Figure 2B**), indicating that MLL2 had only a slight impact on steady state mRNA expression levels early on, which is consistent with previous studies (*30, 33, 34*). It is therefore unlikely that the defects in neuroectoderm differentiation arose from a loss of the primed ESC programme despite the requirement for MLL2 during this period.

Because even small changes in gene expression could have later effects on differentiation, we conducted a detailed analysis of MLL2-dependent changes in gene expression during the transition. With good correlation between log_2_FC changes in the two replicates (**Figure S4A**), we identified 1047 differentially expressed genes (DEGs) with 530 up-regulated and 517 down-regulated genes (p<0.001) (**Figure 2C** and **S4B-C**). To identify direct targets of MLL2, we utilized the published MLL2 ESC ChIP-seq dataset (*35*) to filter for genes that had MLL2 bound within 1 kb of their transcription start site (**Figure S4D**). This revealed that DEGs had a significantly higher likelihood of being MLL2 bound (84% vs 36%; Chi-squared test: p = 10^−267^) (**Figure S4E**) and that there were a significantly higher number of MLL2 peaks within 1 kb of the gene transcription start site (TSS) of these genes (average peak number of 1.1 peaks vs 0.5 peaks; Mann Whitney Test: p = 10^−273^) (**Figure S4F**). It also showed that 90% of MLL2-dependent genes had MLL2 bound: 947 genes out of the 1047 MLL2-dependent DEGs were MLL2-bound. We then separated MLL2-bound DEGs (p<0.001) by cluster and observed cluster-specific changes: more up- than down-DEGs in the 0hr naïve ESC cluster, similar numbers of up- and down-DEGs in the 24hrs clusters and more down- than up-DEGs in the 48-72hrs primed ESC clusters (**Figure 2C and S4B**). To validate these changes, we examined the well-characterised MLL2-target gene *Magohb* (*34, 37*) and found consistent loss of *Magohb* expression in every cluster (**Figure 2D and S4C**). Overall, we conclude that MLL2 was acting directly to modulate transcription at only a few genes during the priming transition, consistent with previous ESC studies (*30, 34, 35*).

Comparison of the *Mll2* cKO DEGs with pathways known to play a role in pluripotency, as defined by PluriNetwork (*50*), highlighted the transcription factors SOX2 and OTX2. OTX2 is required for activation of enhancers during the priming transition (*15, 51*) and SOX2 must be expressed at high levels for activation of neuroectoderm genes (*52*). MLL2 increased the transcriptional amplitude of both genes at 48/72hrs primed ESCs (p = 10^−4^ for *Sox2* at 48hrs A; p = 10^−11^ and p = 10^−9^ for *Otx2* at 48hrs A and at 72hrs respectively) (**Figure 2D**). MLL2 was also bound at the *Sox2* and *Otx2* genes (**Figure S4D**). However, MLL2 only subtly increased transcriptional amplitude at these genes so we took *Sox2* as an example gene to determine whether the small change in *Sox2* mRNA level was enough to cause larger alterations in SOX2 protein expression. Quantitative immunofluorescence revealed that *Mll2* cKO increased SOX2 expression in naïve ESCs (1.3-fold, p < 10^−16^) and decreased it in 72hrs primed ESCs (0.5-fold, p = 10^−14^) (**Figure 2E-F**), comparable to changes observed at the transcript level (**Figure 2D**). Interestingly, SOX2 continued to decrease over time in *Mll2* cKO cells, suggesting that small changes at the mRNA level accumulated over time and were amplified at the protein level.

### MLL2 catalytic activity is not required for neuroectoderm differentiation

Having observed minimal changes in transcription and yet dramatic changes in differentiation, we asked whether MLL2’s methyltransferase activity was playing a role during this priming transition by introducing a BAC transgene into the *Mll2* cKO cell line to express a GFP-tagged MLL2 with an N2650A mutation (MLL2-NCat) that inactivates its catalytic activity (**Figure 3A and S5**). Although we considered generating a homozygous knock-in mutant cell line, confounding factors may arise from ESCs grown for long periods in the absence of MLL2’s catalytic activity which would make data interpretation more difficult. We preferred a conditional approach based on retention of wild-type MLL2 until conditional mutagenesis yields reliance on expression of the N2650A BAC transgene. After tamoxifen induction to ablate endogenous MLL2 expression, immunoprecipitation using anti-GFP antibodies established that the MLL2-NCat expression level was similar to that of the GFP-tagged wild-type MLL2 BAC transgene used to rescue the loss of *Mll2* (**Figure S5A**). ChIP-qPCR of the *Magohb* promoter confirmed that the N2650A mutation impaired catalytic activity (p = 10^−4^) (**Figure S5B**). The validated catalytically-dead MLL2 cell line was successfully differentiated into neural rosettes (**Figure 3B**). Moreover, quantitative immunofluorescence of SOX2 revealed rescue of decreased SOX2 protein level (**Figure 3C-D**). These data establish that MLL2’s catalytic activity is not required for neuroectoderm differentiation. Although the expression of MLL2-NCat was higher than endogenous MLL2 (**Figure S5A**), this does not detract from the conclusion that it could not rescue the catalytic activity of endogenous MLL2 and that this still did not affect differentiation.

**Figure 3.**
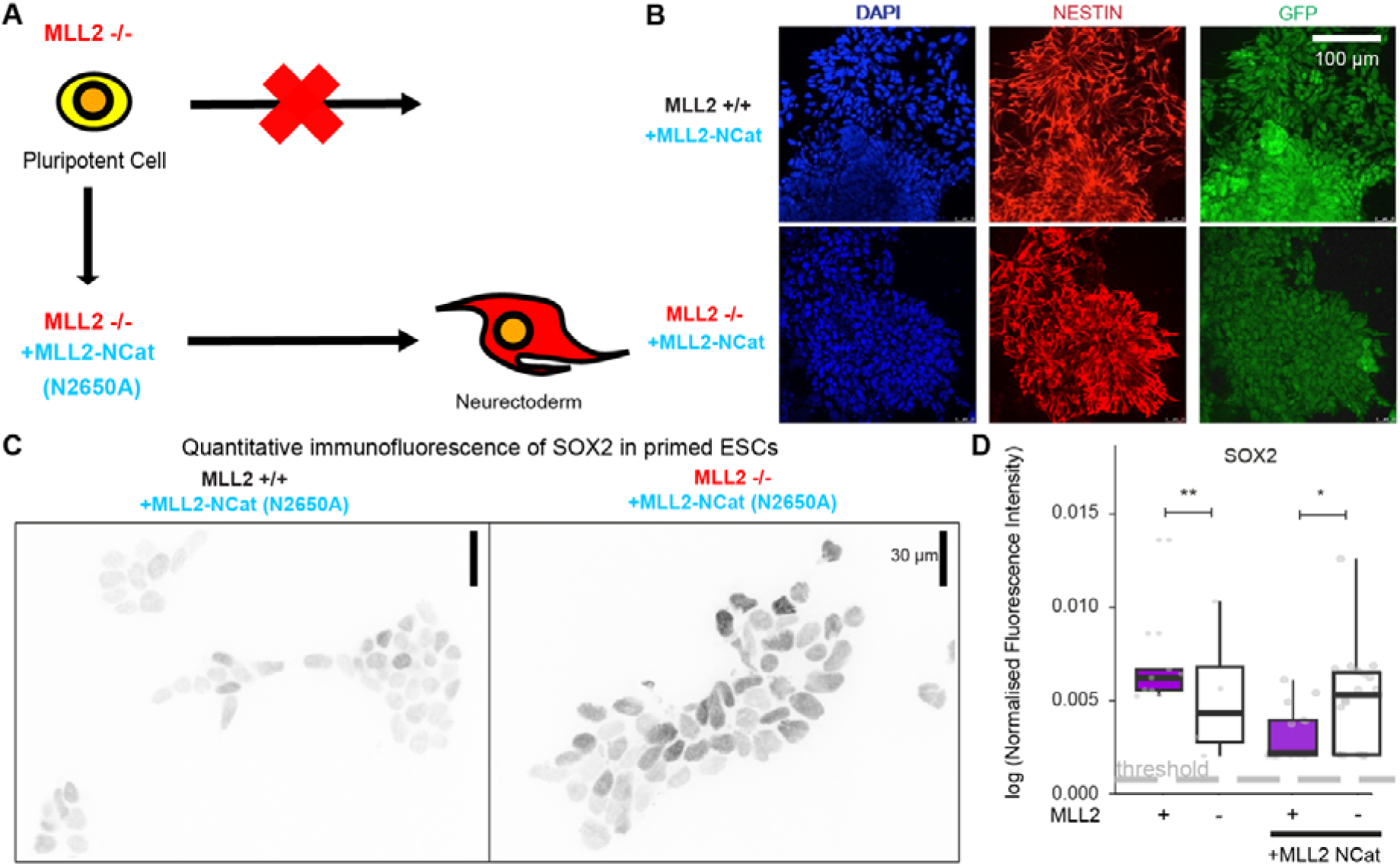
Introduction of catalytically-dead MLL2 (MLL2 NCat) rescues neuroectoderm differentiation and SOX2 expression changes during priming. **(A)** Schematic showing how introduction of a catalytically-dead MLL2 (MLL2 NCat) to *Mll2* conditional knockout (cKO) cells rescues differentiation. **(B)** Fluorescence images of DAPI stain (blue), NESTIN expression (red) and expression of the GFP-tagged MLL2 NCat (green) at the neural rosette stage show that ectopic expression of MLL2 NCat rescues failure of *Mll2* cKO cells to generate neural rosettes (see **Figure 1**). (top) MLL2 +/+ cells ectopically expressing MLL2 NCat. (bottom) MLL2 −/− cells ectopically expressing MLL2 NCat. **(C)** Representative immunofluorescence images of SOX2 in primed ESCs for *Mll2* cKO and control cells with ectopic expression of MLL2 NCat. **(D)** Boxplot of SOX2 expression in primed ESCs for *Mll2* cKO and control cells with/without ectopic expression of MLL2 NCat (images in **Figure 2E and 3C**). Mean fluorescence per cell is shown as grey dots for each field of view and background threshold as a grey dotted line. Number of images (10-100 cells/image): 11/4 (72 hr +/− MLL2), 11/21 (72 hr +/− MLL2 +MLL2NCat-GFP). [unpaired two-sided t-test, **p = 0.007 (72 hr +/− MLL2), *p = 0.03 (72 hr +/− MLL2 +MLL2NCat-GFP).]

### MLL2 knockout has a minor effect on the compartmentalisation of the 3D genome in primed ESCs

Because MLL2 function in neuroectoderm differentiation does not require its methyltransferase activity, we considered other ways MLL2 can influence gene expression, including by decompacting chromatin (through the recruitment of CBP/p300) or by tethering chromatin as suggested by its large size and various chromatin binding domains. To gather evidence of the impact of MLL2 on changes in chromatin other than H3K4 methylation, we monitored 3D genome organisation in wild-type and *Mll2* cKO cells during the priming transition using chromosome conformation capture, which reveals snapshots of the 3D genome through the analysis of DNA sequences that are in physical proximity (*53–62*). We first considered a previous Hi-C chromosome conformation capture study that used the same *Mll2* cKO cells. However, this study (*30*) utilized a culture condition that presents an artificial epigenetic state, termed cycling ESCs, which does not distinguish between naïve and primed states (*4*). We therefore carried out chromosome conformation capture of both naïve and primed ESCs separately. Rather than Hi-C, we applied chromosome conformation capture using Micro-C (*60–62*) to achieve a higher resolution of the 3D genome. Micro-C was carried out in duplicate for wild-type and *Mll2* cKO cells in both naïve and primed ESCs (0 hr and 72 hr), which generated >270 million uniquely mapped reads/sample (**Figure S6A**).

Analysis of genome A-B compartments (*57*) revealed relatively large changes during the priming transition (**Figure 4A**) but smaller changes after the loss of MLL2 (**Figure 4B-C**). Although most of the genome (64%) remained in their original A or B compartments, priming provoked 27% of the genome to switch from A to B compartments with only 9% of the genome switching the other way (**Figure 4A**). There was also a 3.1-fold increase in median B compartment size (p = 10^−8^) relative to the smaller 1.2-fold increase in A compartment size (p = 10^−12^) (**Figure 4D**). In contrast, the loss of MLL2 provoked more subtle changes with few A-B compartment changes in naïve ESCs (**Figure 4B**) but with more changes in primed ESCs with a 12% shift from B to A compartments and only 5% of genomic regions shifting in the opposite direction (**Figure 4C**). This was consistent with a 1.4-fold increase in A compartment size in primed ESCs (p = 10^−4^) with no change in B compartment size (p = 0.4) (**Figure 4D**). We conclude that priming involves a general shift from the A to the B compartment, consistent with DNA methylation and heterochromatinisation known to occur during the priming transition (*63*). MLL2 makes a modest contribution to this shift mainly limited to the expansion of certain B compartments in primed ESCs.

**Figure 4.**
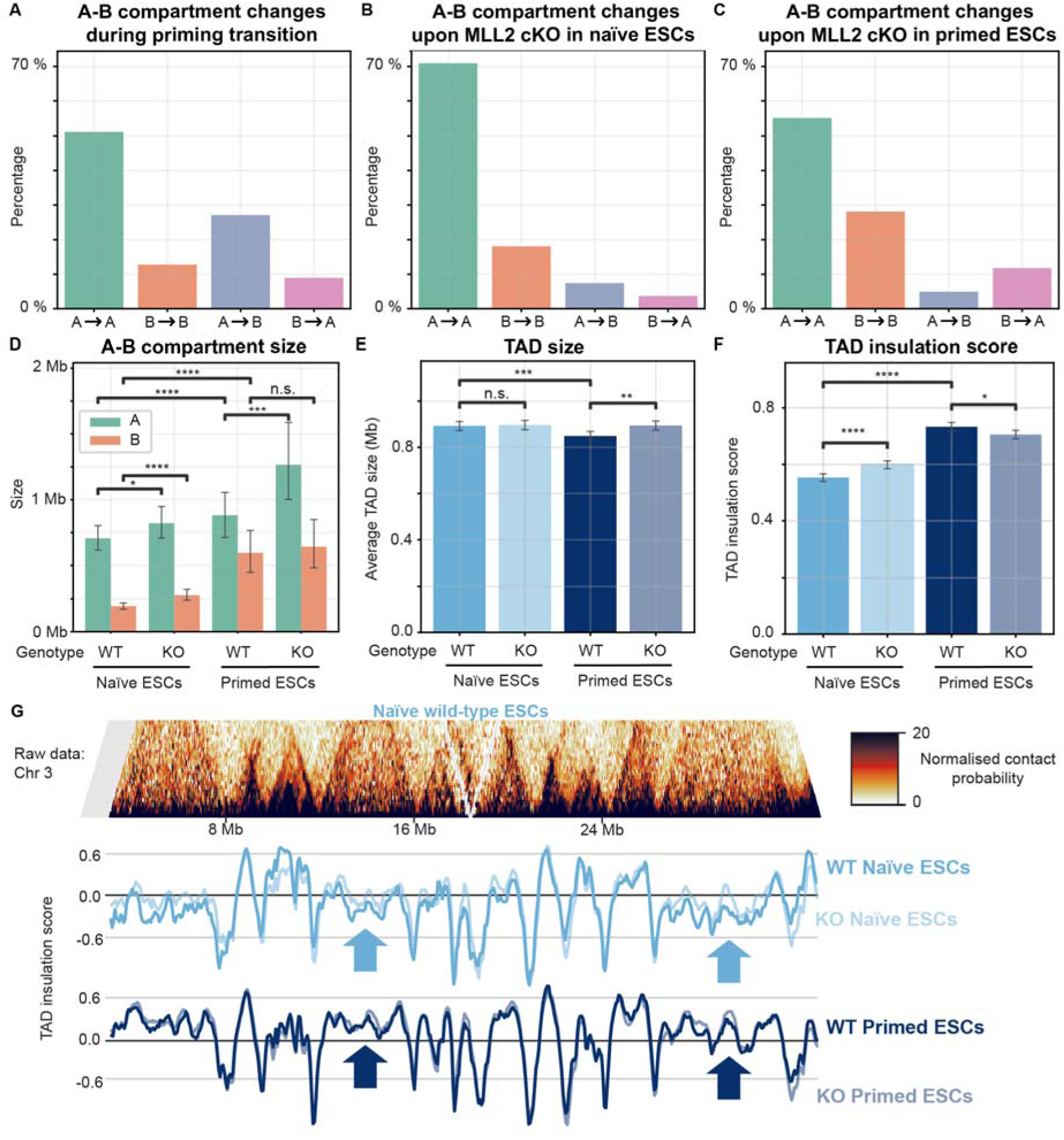
MLL2 knockout has small effect on compartmentalisation of the genome into A-B compartments and TADs (topologically associating domain). **(A)** Percentage of 40 kb genomic bins that transition between A and B compartments when comparing naïve and primed wild-type (WT) ESCs. **(B)** Percentage of 40 kb genomic bins that transition between A and B compartments when comparing WT and *Mll2* cKO (KO) naïve ESCs. **(C)** Percentage of 40 kb genomic bins that transition between A and B compartments when comparing WT and KO primed ESCs. **(D)** Barplot showing mean A/B compartment size in WT and KO conditions for naïve and primed ESCs, with error bars represent 95 % confidence interval. [n = 3031/3010, 2487/2466, 1854/1833 and 1435/1414 A/B compartments identified for WT naïve, KO naïve, WT primed and KO primed ESCs respectively; Mann-Whitney-Wilcoxon two-sided test with Benjamini-Hochberg correction, p = 10^−12^/10^−8^ for WT naïve vs primed ESC A/B compartment size; p = 0.04/10^−12^ for WT vs KO naïve ESC A/B compartment size; p = 10^−4^/0.38 for WT vs KO primed ESC A/B compartment size] **(E)** Barplot of mean TAD size, with error bars represent 95 % confidence interval, shows that KO increases TAD size in primed but not in naïve ESCs. [n = 2831, 2820, 2980 and 2828 TADs identified for WT naïve, KO naïve, WT primed and KO primed ESCs respectively; Mann-Whitney-Wilcoxon two-sided test with Benjamini-Hochberg correction, p = 10^−4^ for WT naïve vs primed ESCs; p = 0.7 for WT vs KO naïve ESCs; p = 10^−3^ for WT vs KO primed ESCs] **(F)** Barplot of mean TAD insulation score, with error bars represent 95 % confidence interval, shows stabilisation of TADs when comparing naïve to primed ESC scores, with more subtle changes observed between WT and KO samples. [n = 2831, 2820, 2980 and 2828 TADs identified for WT naïve, KO naïve, WT primed and KO primed ESCs respectively; Mann-Whitney-Wilcoxon two-sided test with Benjamini-Hochberg correction, p = 10^−76^ for WT naïve vs primed ESCs; p = 10^−8^ for WT vs KO naïve ESCs; p = 0.02 for WT vs KO primed ESCs] **(G)** (Above) Normalised Micro-C contact map for chromosome 3 showing density of contacts allows visual identification of topologically associating domains (TADs). (Below) TAD insulation scores calculated for naïve (light blue) and primed ESCs (dark blue) of priming for WT (darker) and KO (lighter) conditions showing stabilisation of TADs when comparing naïve to primed ESC scores, as indicated by arrows, with more subtle changes observed between WT and KO samples.

Next, we analysed changes at the scale of topologically associating domains (TADs; 500 kb to 3 Mb). Based on the TAD insulation score, which reflects the likelihood of interactions within a TAD versus outside of it, we found that priming provoked larger changes than the loss of MLL2 (**Figure 4E-G**). Specifically, we observed a decrease in TAD size during the priming transition (p = 10^−4^). Intriguingly, this decrease that was not observed in primed ESCs upon loss of MLL2 (p = 10^−3^) (**Figure 4E**). We also observed an increase in TAD formation during priming, with insulation scores increasing from 0.553 to 0.733 (p = 10^−76^) (**Figure 4F**). Loss of MLL2 related to only minor changes in this score, with a slight increase from 0.553 to 0.599 in naïve ESCs (p = 10^−8^) and a slight decrease from 0.733 to 0.706 in primed ESCs (p = 0.02). Therefore, in agreement with the conclusion drawn by Mas et al. (2018) using cycling ESCs, MLL2 has only minor effects on TAD organization in naïve and primed ESCs, although MLL2 was required to decrease TAD size during the priming transition.

### MLL2 knockout influences 3D chromatin looping

In the replicates, global comparisons of A-B compartments and TADs were highly correlated (**Figure S6B**). Therefore, we compiled replicates to increase the chromatin looping resolution, which allowed robust analysis at 10kb resolution. Previous studies indicated that priming involves changes in long-range interactions (> 1Mb) (*22, 23*) so 10kb resolution is more than sufficient to detect these changes. High-confidence 3D chromatin loops were mapped using FitHiC2 with a false discovery rate (FDR) of 0.05 (*64*). Because FitHiC2 does not normalise against read-depth, we downsampled to ensure similar numbers of paired reads for our comparisons. In wild-type conditions, we observed a 2.0-fold increase in genome-wide high-confidence loops detected during the transition from naïve to primed ESCs (13,137 vs 26,059 loops for naïve vs primed ESCs respectively) (**Figure 5A and S6A**). Loss of *Mll2* led to both a gain and loss of loops in naïve and primed ESCs (**Figure 5B-C**). While this led to a small but significant decrease in overall loop size in naïve ESCs, it led to a small increase in loop size in primed ESCs (p = 10^−6^ and 10^−6^ for naïve and primed ESCs respectively) (**Figure 5D**). We concluded that loss of *Mll2* leads to observable rewiring of chromatin loops and changes in loop size, with differences between naïve and primed ESCs.

**Figure 5.**
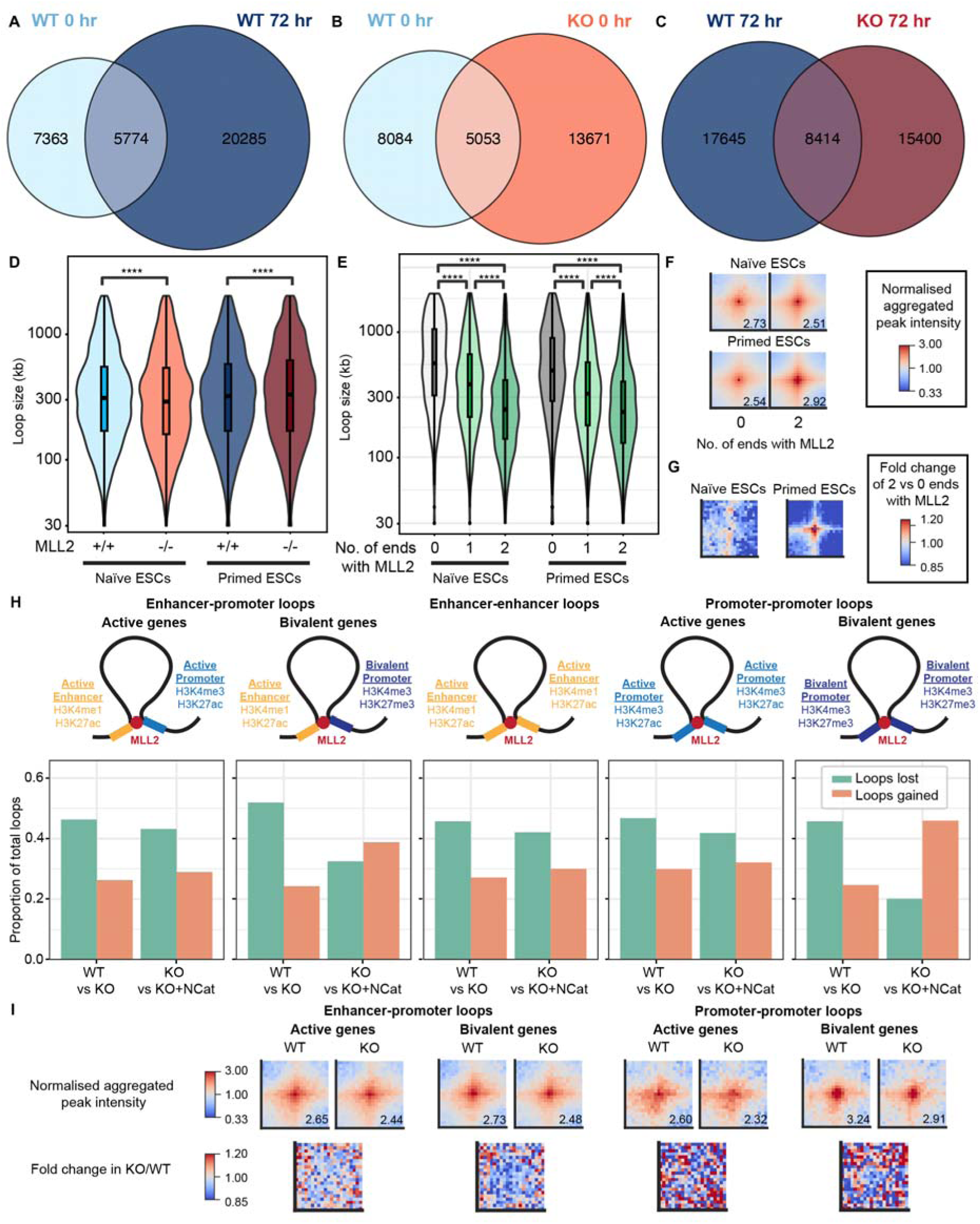
MLL2 knockout rewires 3D chromatin looping but MLL2 does not require its catalytic activity to form enhancer-promoter loops at bivalent genes in primed pluripotent cells. **(A)** Euler diagram showing overlap of loops detected in naïve ESCs (0 hr) vs in primed ESCs (72 hr). **(B)** Euler diagram showing overlap of loops detected in WT vs KO cells for naïve ESCs (0 hr). **(C)** Euler diagram showing overlap of loops detected in WT vs KO cells for primed ESCs (72 hr). **(D)** Box plot showing loop size distributions in WT vs KO cells for naïve and primed ESCs. [Mann-Whitney-Wilcoxon two-sided test, p = 10^−6^ for WT vs KO naïve ESCs; p = 10^−6^ for WT vs KO primed ESCs] **(E)** Box plot showing distribution of loop sizes in WT cells for loops without MLL2, with MLL2 bound at one of the loop anchors or with MLL2 bound at both of the loop anchors for naïve and primed ESCs [Mann-Whitney-Wilcoxon two-sided t-test, p = 10^−58^, 10^−156^ and 10^−238^ for MLL2 at 0 vs 1 end, 1 vs 2 ends and 0 vs 2 ends in naïve ESCs; p = 10^−204^, 10^−157^ and 10^−539^ for MLL2 at 0 vs 1 end, 1 vs 2 ends and 0 vs 2 ends in primed ESCs] **(F)** Aggregate peak analysis shows that loops with MLL2 on both ends have lower looping strength than loops without MLL2 in naïve ESCs but higher looping strength in primed ESCs. Observed/expected peak intensity score normalised by corner pixels shown for 100 kb region centred on loops (FDR<0.05). Peak intensity score of loop anchor (middle pixel) is indicated. **(G)** Fold-change in peak intensity score for KO/WT values shown in (F). **(H)** (Row 1) Schematic showing how categories of loops are defined as enhancer-promoter loops at active or bivalent genes, enhancer-enhancer loops and promoter-promoter loops at active or bivalent genes. (Row 2) Bar plots for categories in Row 1 showing proportion of loops gained (green) or lost (orange) when comparing WT and KO primed ESCs but also when comparing KO to KO primed ESCs that express the N2650A catalytically-dead mutant form of MLL2 (KO; KO+NCat). [Fisher’s exact test with false discovery rate (FDR) multiple test correction, p-values: 10^−21^/10^−9^, 10^−23^/0.04, 10^−16^/10^−6^, 10^−5^/0.04 and 10^−6^/10^−8^ (E-P, E-BP, E-E, P-P and BP-BP loops in WT vs KO/ KO vs KO+NCat)]. **(I)** Aggregate peak analysis for categories in (G) when comparing WT and KO primed ESCs. (Top) Observed/expected peak intensity score shown for 100 kb region centred on wild-type loops (FDR<0.05) and normalised by corner pixels. Peak intensity score of loop anchor (middle pixel) is indicated. (Bottom) Fold-change in peak intensity score for KO/WT values shown above.

To identify changes related to MLL2 binding, we used a published MLL2 ChIP-seq dataset (*35*) to subset Micro-C loops with (i) no MLL2 sites at the anchors, (ii) MLL2 on one end of the loop and (iii) MLL2 at both ends of the loop (within our 10kb resolution). This revealed 11,574 MLL2-associated loops in naïve ESCs (88% of total loops) and 19,718 loops in primed ESCs (76% of total loops), indicating a 1.7-fold increase in the number of MLL2-associated loops during the priming transition. We next characterised the size and strength of these MLL2-associated loops. Loops with MLL2 binding sites at both ends were smaller than loops with MLL2 at only one end or at neither end (**Figure 5E**). Moreover, aggregate peak analysis of loops with and without MLL2, revealed that, although MLL2-associated loops (with MLL2 at one or both ends) were weaker than loops without MLL2 in naïve ESCs, this trend changed during the priming transition with MLL2-associated loops now stronger than loops without MLL2 (**Figure 5F-G**). We concluded from this that (i) MLL2 is associated with a large proportion of loops in both naïve and primed ESCs; (ii) the number of MLL2-associated loops increases during the priming transition; (iii) MLL2 is associated with smaller loops; and (iv) MLL2 has an inverse effect on the strength of loops in naïve versus primed ESCs.

### MLL2 contributes in a non-catalytic manner to loops involving bivalent promoters

MLL2 is bound at bivalent promoters in ESCs, and its loss reduces H3K4me3 levels at these promoters (*35*). However, MLL2 is also bound to many other Pol II promoters where its loss has no impact on H3K4me3 levels (*35*). This evidence for a special relationship between MLL2 and bivalent promoters led us to first focus on loops involving bivalent promoters. As noted in several past papers, H3K27me3 levels are substantially lower in naïve ESCs (*4, 16*). Hence there are far fewer bivalent promoters in naïve ESCs with bivalent promoters arising primarily during the transition to primed ESCs (**Figure S7A**). Therefore, we initially focused our analysis on loops and bivalent promoters in primed ESCs. Using published ChIP-seq datasets (*16*), we identified the correspondence between loop anchors and active enhancers, active promoters or bivalent promoters to classify loops into 5 main categories (bivalent promoter-bivalent promoter; enhancer-bivalent promoter; enhancer-enhancer; enhancer-promoter; promoter-promoter). In primed ESCs, we observed 168 loops between bivalent promoters and more than half (102 loops) were lost in the absence of *Mll2* (p = 10^−6^) although we did observe 55 de novo loops in knockout cells (**Figure 5H and S7B**). Similar trends were observed even when a more stringent FDR was chosen (q<10^−5^) (**Figure S7B**). Aggregate peak analysis also followed the overall trend showing a loss in looping strength between bivalent promoters (**Figure 5I**). Intriguingly, Micro-C of the *Mll2-NCat* cell line (**Figure S6A**) revealed that most of these loops were restored by *Mll2-NCat* (p = 10^−8^) (**Figure 5H and S7B**), albeit with some variance, since the specific promoter-promoter loops being restored were not always the same as in wild-type conditions (**Figure S7C**). We considered whether the loss of looping at MLL2 target sites might reflect decreased MNase accessibility in the absence of MLL2. However, Micro-C datasets were ICE-normalized to account for systematic biases including differences in cutting efficiency between samples. Moreover, MNase digestion efficiency (assessed through percentage of mononucleosomes after digestion) was comparable in *Mll2* cKO cells compared to WT (57 9 % vs 68 3 % mononucleosomes in WT vs cKO naïve ESCs and 66 10 % vs 64 5 % mononucleosomes in WT vs cKO primed ESCs), arguing against reduced cutting as an explanation.

In contrast, although loops in primed ESCs between active elements (enhancer-enhancer, enhancer-promoter and promoter-promoter) were also affected by the loss of *Mll2* (p = 10^−16^, 10^−21^ and 10^−5^), these losses were not rescued by *Mll2-NCat* (**Figure 5H-I and S7B**), suggesting that active elements require the H3K4me3 mark for enhancer-promoter looping. Notably, loops between active enhancers and bivalent promoters showed an intermediate effect, impacted by the loss of *Mll2* (p = 10^−23^) but only partially rescued by *Mll2-NCat* (**Figure 5H and S7B**). Loss of *Mll2* also promoted a decrease in loops between non-specific/undefined genomic regions and enhancers, active promoters or bivalent promoters in primed ESCs (p = 10^−40^, 10^−39^ and 10^−126^) and these loops were not rescued by *Mll2-NCat* (**Figure S8**), suggesting that MLL2 indirectly increases the formation of additional undefined loops at enhancers and promoters. Together with the impact of MLL2 on looping, loop strength and size, the relationship between MLL2 and loops involving bivalent promoters strengthens the conclusion that MLL2 directly contributes to looping at bivalent promoters, with gene ontology showing that these loops are linked to neuroectoderm-specifying genes (**Figure S7D**), while suggesting a role for MLL2’s catalytic activity in the indirect formation/maintenance of loops at active enhancers/promoters.

### MLL2 knockout weakens chromatin loops associated with gene activation while enhancing loops associated with DNA methylation and heterochromatin in primed ESCs

Because loss of *Mll2* led to a decrease in enhancer/promoter looping but also an increase in non-specific looping, we further characterised loops mediated/inhibited by MLL2 by taking advantage of published ChIP-seq datasets (*65*) to identify chromatin binding proteins enriched at the loop anchor regions gained/lost in *Mll2* cKO cells.

In naïve ESCs, *Mll2* cKO led to a loss of loops enriched for the Polycomb complex PRC1 (CBX2, CBX6, RYBP) (**Figure 6A and 6C**). This initially appeared surprising given the antagonism between MLL2 and Polycomb (*30, 36*). However, because MLL2 and Polycomb co-bind, for example at bivalent promoters, it is likely that they act together to increase looping at these co-bound sites. Loops lost also included proteins involved in enhancer-promoter communication in naïve ESCs such as the transcription factor KLF4, the Mediator complex (MED26) and the TFIID complex (TBP). At the same time, *Mll2* cKO led to more loops enriched for the NuRD complex (MBD3, MTA2, SALL4). H3K4me3 marks recruit the TFIID complex and block NuRD complex recruitment (*66*) so recruitment of these proteins may be linked to MLL2’s catalytic activity.

**Figure 6.**
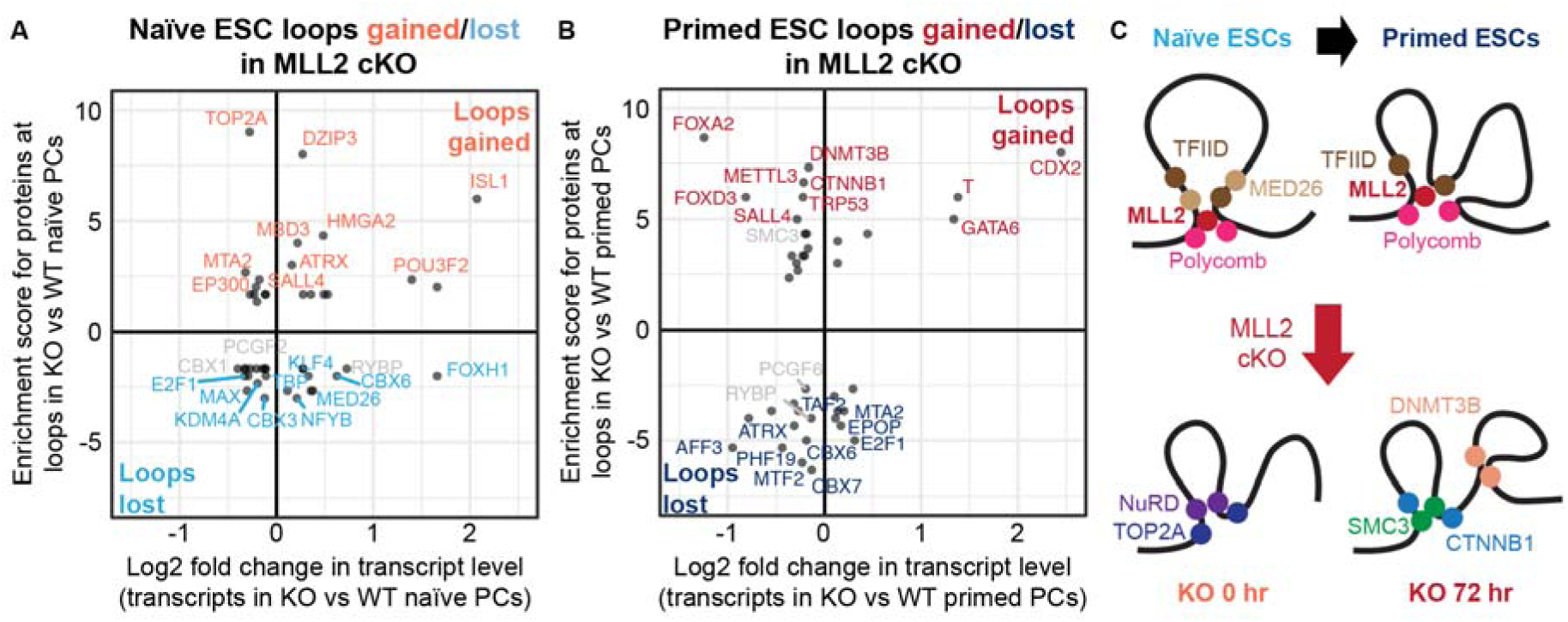
MLL2 knockout weakens 3D chromatin loops linked to gene activation or bivalent genes and enhances loops linked to DNA methylation and heterochromatin in primed ESCs. **(A)** Correlation plot showing how changes in protein enrichment at loops relate to change in expression of enriched proteins when comparing *Mll2* cKO (KO) (top right) to wild-type (bottom left) naïve ESCs (0 hr). Proteins enriched in wild-type and KO ESCs are coloured light blue and light red respectively. **(B)** Correlation plot showing how changes in protein enrichment at loops relate to change in expression of enriched proteins when comparing KO (top right) to wild-type (bottom left) primed ESCs (72 hr). Proteins enriched in wild-type and *Mll2* cKO ESCs are coloured dark blue and dark red respectively. **(C)** Schematic showing changes in proteins at loops during priming of pluripotent cells for wild-type and *Mll2* cKO cells.

In primed ESCs, *Mll2* cKO again led to a loss of loops enriched for Polycomb complexes PRC1 and PRC2 (CBX7, CBX6, PCGF6, RYBP) (**Figure 6B-C**). At the same time, we observed more loops enriched for DNMT3B, consistent with MLL2’s known ability to block DNA methylation (*34, 36, 37*). Interestingly, we also observed more loops for the cohesin complex (SMC3), suggesting that MLL2 blocks loop extrusion, for proteins linked to the WNT signalling pathways (β-catenin), consistent with a link between MLL proteins and the WNT signalling pathway (*67, 68*) and for p53, consistent with increased apoptosis upon MLL2 knockout (*33*).

We concluded from our analysis that MLL2 likely acts both directly, supported by reduced chromatin looping of MLL2-bound regions, but also indirectly with increased chromatin looping of DMT3B-, Polycomb- and cohesin-bound regions.

### MLL2 tethers chromatin genome-wide during the priming transition

Because MLL2 substantially influences looping in ESCs, we asked whether it modulates chromatin tethering on a genome-wide scale by using 3D single-molecule localisation microscopy (SMLM) to track the mobility of individual nucleosomes within the nucleus +/− MLL2 (**Figure 7A**). SMLM has previously revealed that chromatin mobility decreases in the presence of cohesin loops and increases upon histone acetylation or in the presence of specific chromatin remodellers (*69–72*). It has also shown that chromatin mobility depends on cell type, with differences observed between ESCs and more differentiated cell types (*73*).

**Figure 7.**
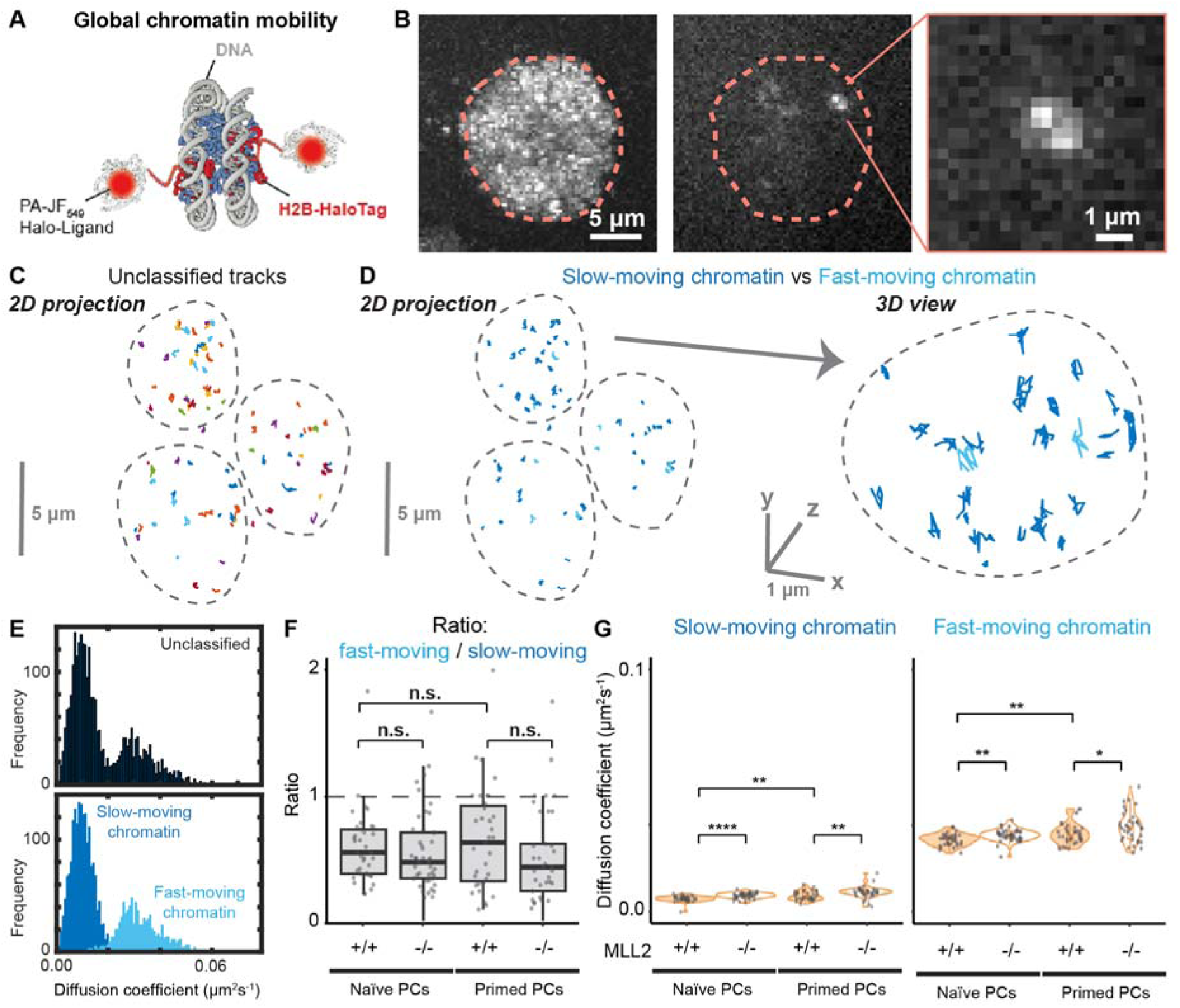
MLL2 knockout increases chromatin mobility genome-wide during priming of pluripotent cells. **(A)** Schematic showing PA-JF_549_-HaloTag-tagged histone H2B. **(B)** (Left) Sum projection of single-molecule localisation microscopy (SMLM) video shows nuclear localisation of histone H2B within a nucleus indicated by an orange dotted line. (Middle) Representative SMLM image showing single PA-JF_549_-HaloTag-tagged histone H2B molecules in a live embryonic stem cell (ESC) imaged using a double helix point spread function (DHPSF) microscope. Each molecule appears as two dots where the midpoint encodes the x,y position and where the angle between the dots encodes the z position. The nucleus is indicated by an orange dotted line based on the sum projection (Right) Inset showing zoomed in image of a single histone H2B molecule. **(C)** Tracks pre-classification. **(D)** (Left) Tracks classified as slow- (dark blue) and fast-moving chromatin (light blue). (Right) Inset showing zoomed in image of a few classified trajectories. **(E)** Histogram showing apparent diffusion coefficient D_app_ of histone H2B trajectories (above) pre-classification and (below) post-classification into slow- or fast-moving chromatin. **(F)** Box-and-violin plot showing ratio of H2B trajectories classified as fast-moving/slow-moving for naïve and primed ESCs in WT/KO conditions. Dots represent ratios calculated from independent videos (n>36 videos, 3-5 cells per video) [unpaired two-sided t-test, p-values: 0.8 (WT naïve vs primed ESCs), 0.3 (WT vs KO in naïve ESCs), 0.09 (WT vs KO in primed ESCs)] **(G)** Box-and-violin plot showing apparent diffusion coefficient D_app_ of (left) slow- and (right) fast-moving chromatin for naïve and primed ESCs in WT/KO conditions. Dots represent mean values calculated from independent videos (n>36 videos, 3-5 cells per video) [unpaired two-sided t-test, p-values: **10^−3^ (slow-moving WT naïve vs primed ESCs), ****10^−6^ (slow-moving WT vs KO in naïve ESCs), **10^−3^ (slow-moving WT vs KO in primed ESCs), **10^−3^ (fast-moving WT naïve vs primed ESCs), **10^−3^ (fast-moving WT vs KO in naïve ESCs), *0.02 (fast-moving WT vs KO in primed ESCs)]

To conduct SMLM, we introduced Halo-tagged histone H2B into the *Mll2* cKO cell line, validating expression and nuclear localisation through labelling of cells with a JFX_549_ dye (**Figure 7B and S9A**). We then labelled H2B for SMLM using the photo-activatable PA-JF_549_ dye (*74*). Cells were imaged using highly inclined and laminated optical (HILO) illumination and double-helix point spread function microscopy (*71, 75*). This allowed us to track individual nucleosomes in 3D through a 4 μm slice of the nucleus (**Figure 7 and S9A**). Individual PA-JF_549_-HaloTag-tagged histone H2B molecules were detected as evidenced by single-step photobleaching profiles (**Figure S9B-C**). We collected videos at 500 ms temporal resolution (5 fields of view per replicate, 3 replicates, 2000 frames) because 500 ms exposures detect nucleosomal motion while motion blurring any freely diffusing H2B (*71, 76*). Using fixed cell datasets, we determined the precision at which we could localise a single H2B molecule (54 nm 10 nm in x, 68 nm 8 nm in y and 134 nm 8 nm in z) and found that H2B displacements were significantly higher in all live cell datasets when compared to fixed cell datasets (**Figure S9D**).

To analyse these H2B trajectories, we extracted biophysical parameters from sub-trajectories of 7 frames (3.5 s) to determine how fast chromatin diffuses – the apparent diffusion coefficient D_app_. Chromatin has viscoelastic properties, so its mobility depends on the time and length scale used for measurements (*77*). We chose this timescale because it corresponds to a length scale of ∼250 nm. This corresponded to the size of TADs (*70, 78, 79*), allowing us to test changes in TAD size we had previously observed (**Figure 4E-G**) but also corresponded to the size of MLL2-associated loops (**Figure 5D-E**). Also, at this time and length scale, chromatin mobility does not change during the cell cycle (*80*), whereas cell cycle effects have been observed at timescales of a minute or longer (*81*). As previously described, this revealed slow- and fast-moving sub-diffusive chromatin states (**Figure 7C-E**) (*71, 72, 82, 83*). Although the nature of slow- and fast-moving chromatin remains unresolved, meaningful information can still be extracted regarding whether perturbations increase tethering or chromatin compaction (lower D_app_) (*69, 71, 72*). We therefore took advantage of our published 4-parameter classification algorithm to classify individual trajectories as either slow- or fast-moving prior to extraction of D_app_ (**Figure 7C-E**) (*71*), and then used these parameters to determine how MLL2 influences slow- or fast-moving chromatin during priming.

We first determined how chromatin mobility changes as cells transition from naïve to primed ESCs. There was no change in the proportion of H2B molecules observed in slow- and fast-moving chromatin states (p = 0.8) (**Figure 7F and S9E**). However, an increase in D_app_ during the priming transition was apparent for both the slow- and fast-moving chromatin states (p = 10^−3^ for both) (**Figure 7G**), indicating that chromatin mobility was increased. We next asked whether similar changes were occurring in the absence of *Mll2*. Although no change in the proportion of H2B molecules in slow- and fast-moving chromatin states was apparent (p = 0.3 and 0.09 for naïve and primed ESCs) (**Figure 7F**), loss of *Mll2* increased D_app_ for slow- and fast-moving chromatin states in both naïve and primed ESCs (p = 10^−6^ and 10^−3^ for naïve ESCs; p = 10^−3^ and 0.02 for primed ESCs) (**Figure 7G**). We concluded from this that MLL2 restricts chromatin mobility in both naïve and primed ESCs, consistent with a role in chromatin tethering, although the effect was larger in primed ESCs possibly because there was an overall increase in chromatin mobility during the naïve to primed transition. This was consistent with the observed increase in TAD and loop size observed upon loss of *Mll2*, particularly in primed ESCs (**Figure 4E and 5D**).

Finally, since enhancer-promoter looping is partially rescued by *Mll2-NCat*, we introduced Halo-tagged histone H2B into the *Mll2-NCat* cell line (**Figure S5D**) We again saw no change in the proportion of H2B molecules observed in slow- and fast-moving chromatin states (p = 0.14 and 1.0 for naïve and primed ESCs) (**Figure S9H**). However, in contrast to the *Mll2* cKO cell line, we no longer observed an increased D_app_ for the fast-moving chromatin state (p = 0.5 and 0.7 for naïve and primed ESCs) (**Figure S9I**). For slow-moving chromatin, we observed a minor increase in D_app_ in primed ESCs (p = 0.01) but no change in naïve ESCs (p = 0.5). Consequently, we concluded that *Mll2-NCat* partially, although incompletely, restored the chromatin mobility changes observed after loss of endogenous *Mll2*.

### MLL2 exhibits cell type-specific effect on tethering of enhancer-promoter loop at *Sox2* gene during priming transition

Having observed that MLL2 tethers chromatin genome-wide, we next focussed on how it influences looping at specific genes. For gene-specific analysis, we applied more stringent conditions only visualising loops that had a FDR<10^−5^. This revealed changes in chromatin looping at our previously identified DEGs *Sox2* and *Otx2* (**Figure 8A-B** and **S10**).

**Figure 8.**
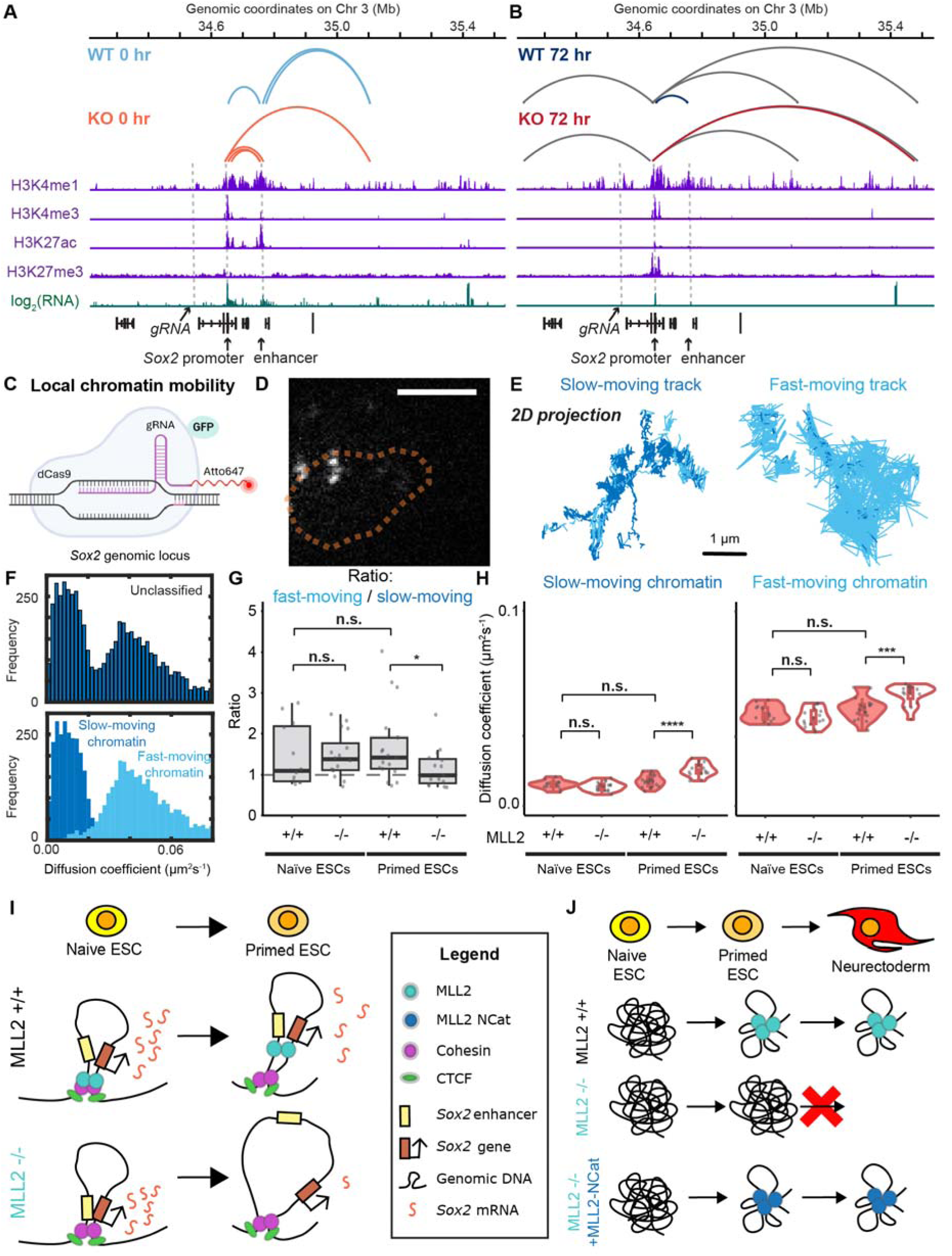
MODEL: MLL2 knockout untethers enhancer-promoter loop at *Mll2*-dependent gene *Sox2* during priming of pluripotent cells, disrupting *Sox2* expression levels and neuroectoderm differentiation. **(A-B)** High-confidence loops (FDR<10^−5^) alongside published RNA-seq and ChIP-seq of histone marks (*16*) at *Sox2* gene in **(A)** naïve and **(B)** primed ESCs for wild-type (WT) and *Mll2* cKO cells (KO). Grey dotted line represents *Sox2* enhancer and promoter. Grey loops are common to both conditions, light blue to WT naïve ESCs, light red to KO naïve ESCs, dark blue to WT primed ESCs and dark red to KO primed ESCs. **(C)** Schematic showing dCas9-GFP targeted to genomic repeat near the *Sox2* gene locus using a specific Atto647-tagged gRNA. **(D)** Representative SMLM image of three labelled *Sox2* gene loci in a live ESC imaged using a DHPSF microscope. The nucleus is indicated by an orange dotted line based on the bright field image. **(E)** Representative trajectories of *Sox2* gene loci that are either mostly slow-moving or mostly fast-moving. **(F)** Histogram showing apparent diffusion coefficient D_app_ of *Sox2* gene sub-trajectories (above) pre-classification and (below) post-classification into slow- or fast-moving chromatin. **(G)** Box-and-violin plot showing ratio of *Sox2* gene sub-trajectories classified as fast-moving/slow-moving gene for naïve and primed ESCs in WT/KO conditions. Dots represent ratios calculated from independent videos (n>12 videos, 3-5 cells per video) [unpaired two-sided t-test, p-values: 0.3 (WT naïve vs primed ESCs), 1.0 (WT vs KO in naïve ESCs), *0.03 (WT vs KO in primed ESCs)] **(H)** Box-and-violin plot showing apparent diffusion coefficient D_app_ of (left) slow- and (right) fast-moving *Sox2* gene for naïve and primed ESCs in WT/KO conditions. Dots represent mean values calculated from independent videos (n>12 videos, 3-5 cells per video) [unpaired two-sided t-test, p-values: 0.08 (slow-moving WT naïve vs primed ESCs), 0.2 (slow-moving WT vs KO in naïve ESCs), ****10^−5^ (slow-moving WT vs KO in primed ESCs), 0.2 (fast-moving WT naïve vs primed ESCs), 0.2 (fast-moving WT vs KO in naïve ESCs), ***10^−4^ (fast-moving WT vs KO in primed ESCs)]. **(I)** Schematic showing enhancer-promoter looping and transcriptional changes at *Sox2* gene during priming transition. In this transition, the CTCF binding pattern changes leading to a larger chromatin loop at the *Sox2* locus. Throughout priming MLL2 tethers a short enhancer-promoter *Sox2* loop. This increases enhancer-promoter spatial proximity and contact frequency, maintaining *Sox2* transcription in primed ESC within the larger chromatin loop. When MLL2 is lost in naive ESC, a short-range CTCF chromatin loop maintains the *Sox2* promoter-enhancer proximity and high *Sox2* transcription. When MLL2 is lost in primed ESC, the larger CTCF chromatin loop is not able to maintain close *Sox2* promoter-enhancer spatial proximity, causing lower contact frequency and transcriptional activity. **(J)** Schematic showing how in wild-type cells MLL2 increases tethering of enhancer-promoter loops at bivalent genes during priming of pluripotent cells to allow neuroectoderm differentiation. *Mll2* cKO disrupts enhancer-promoter loops and increases formation of repressive loops during priming of pluripotent cells, blocking neuroectoderm differentiation. Introduction of a catalytically-dead MLL2 restores enhancer-promoter loops at bivalent genes and restores differentiation.

We became particularly interested in the *Sox2* gene because it transitions from an active to a bivalent promoter during the priming transition (**Figure 8A-B**) and because loss of *Mll2* led to cell-type specific effects, upregulating *Sox2* in naïve ESCs while downregulating it in primed ESCs (**Figure 2D-F**). Given the strong connection between MLL2 and bivalent genes, we hypothesised that loss of *Mll2* would have a different effect on looping as *Sox2* transitions from an active in naïve ESCs to a bivalent promoter in primed ESCs. First, to validated our looping analysis, we confirmed that, in naïve ESCs, *Sox2* does indeed form a well-characterised enhancer-promoter loop that has previously been shown to be driven by CTCF/SMC3-driven loop extrusion (**Figure 8A**) (*84*). Interestingly, while loss of *Mll2* had minimal impact on looping between this active enhancer and promoter in naïve ESCs (if anything, there are more loops called in the knockout), it led to a loss of looping between the active enhancer and bivalent promoter in primed ESCs (**Figure 8A-B**). This was consistent with the transcriptional changes observed (**Figure 2D-F**) and suggested that MLL2 maintains *Sox2*’s enhancer-promoter loop as it transitions from an active to a bivalent promoter during the priming transition.

To confirm these looping changes at the *Sox2* gene, we assessed local chromatin mobility at the *Sox2* promoter (**Figure 8C**). We introduced GFP-tagged catalytically-dead dCas9 into the *Mll2* cKO cell line alongside an Atto647-labelled guide RNA sequence that was designed to target a 14-repeat genomic region 110 kb upstream of the promoter of *Sox2*, a region with no functional histone marks or enhancers that influence *Sox2* transcription and in a direction opposite to the *Sox2* enhancer-promoter loop (**Figure 8A-C, S4D and S9F**). We then repeated 3D SMLM and classified trajectories as slow- or fast-moving as described above for H2B (**Figure 8D-F and S9G**). This revealed that the *Sox2* gene was more likely to be observed in the fast-moving chromatin state than previously observed genome-wide using H2B (**Figure 8G**). We also found that loss of *Mll2* reduced the proportion of *Sox2* gene trajectories classified as fast-moving but only in primed ESCs (p = 0.03) (**Figure 8G**). Interestingly, consistent with our H2B findings, loss of *Mll2* again increased D_app_ for both slow- and fast-moving chromatin states but this time only in primed ESCs (p = 0.2 and 0.2 for naïve ESCs; p = 10^−5^ and 10^−4^ for primed ESCs) (**Figure 8H**). Notably, these changes are unlikely to be primarily driven by transcription because no changes in chromatin mobility are observed during priming despite transcriptional down-regulation of *Sox2* (**Figure 2D-F**). Moreover, D_app_ was calculated at timescales below 10 seconds because transcription does not influence the D_app_ of the *Sox2* promoter at these timescales (*84*). We concluded from this that, although MLL2 broadly restricts chromatin mobility across the genome, its impact on specific genes is likely influenced by cell state-specific chromatin loops present at these genes.

## Discussion

H3K4 methylation is important for eukaryotic promoter activity, and epigenetic regulation. This has focused attention on the enzymatic activity of the MLLs to the distraction from their other remarkable characteristics. Indeed, evidence that the H3K4 methyltransferase activities are not required for MLL functions was unexpected (*29, 31, 85*) and is still largely ignored. Like the *Drosophila* orthologs Trithorax and Trr/Lpt (Trithorax-related/Lost PHDs of Trr), the MLLs are among the largest nucleoplasmic proteins, but their highly conserved SET H3K4 methyltransferase domains comprise less than 10% of the protein. Here, we describe how other regions of MLL2, which include extensive IDRs and multiple chromatin interaction modules, convey an essential function during neuroectodermal differentiation.

Using conditional mutagenesis, we found that the requirement for MLL2 in neuroectodermal differentiation is exerted many cell cycles before differentiation is impaired and is independent of its H3K4 methyltransferase activity (**Figure 1 and 3)**. Consistent with findings that none of the MLLs are required in ESCs or for mouse development before implantation (Ernst et al., 2004, Glaser et al., 2009; Ashokkumar et al., 2020), scRNA-seq analysis confirmed that MLL2 is not required for the naïve to primed ESC transition (**Figure 2**). During the transition, the impact of subtle reductions in gene expression, including reduced expression of the important neuroectodermal transcription factors, *Sox2* and *Otx2*, could explain the downstream defects, because high SOX2 expression is required for differentiation (*52*).

To link the non-enzymatic functions of MLL2 to changes in gene expression, we used Micro-C to map 3D genome organisation in the naïve to primed ESC transition. Only limited associations between MLL2 and A/B compartments or TADs were observed. In contrast, there was substantial loss of 3D enhancer-promoter looping, particularly at bivalent genes related to neuroectoderm differentiation (**Figure 4-5**). Notably, we found that, while MLL2’s catalytic activity had some role in enhancer-promoter looping at active genes, MLL2 did not require its catalytic activity to stabilise enhancer-promoter loops near these bivalent genes. We therefore tracked chromatin mobility using SMLM to test whether MLL2 stabilises 3D chromatin loops. This revealed that it does indeed restrain chromatin mobility both globally and at the MLL2-bound *Sox2* gene locus (**Figure 7-8**), consistent with this function. Consequently, we propose that MLL2 stabilises 3D enhancer-promoter loops at bivalent genes through its chromatin binding domains rather than its enzymatic activity, and that this function is required for neuroectoderm differentiation (**Figure 8I-J and S7D**).

Our proposition relies in part on correlating loops mapped by Micro-C, which like other Hi-C methods is far from quantitative, with genome and chromatin features. The strongest relationship that emerged from these correlations identified loops that are lost in the absence of MLL2 (which we term ‘MLL2 loops’) with nearby bivalent promoters (**Figure 5H-I**). The concordance between MLL2 and bivalent promoters had previously been identified based on H3K4me3 ChIP-sequencing after MLL2 conditional mutagenesis and RNAi knock-down (*35, 86*). Here, the independent identification of a relationship between MLL2 and bivalent promoters by Micro-C analysis lends confidence to the qualities of our data and analysis. Further confidence derives from the prominent relationship between MLL2 loops and Polycomb binding sites (**Figure 6**), which resonates with the well-established interactions between Trithorax and the Polycomb-Group (*87–89*). Other associations between MLL2 loops and cohesin, TFIID or β-catenin warrant further investigation.

Understanding how MLL2-dependent changes in chromatin organisation or mobility influence transcription is not fully clear from our studies because MLL2 apparently secures enhancer-promoter loops at bivalent genes prior to transcriptional activation. Nevertheless, some insight can be gained from specific cases such as *Otx2* or *Sox2*, both of which acquire H3K27me3 to become bivalent promoters during the priming transition. Loss of *Mll2* results in loss of enhancer-promoter loops at both genes in primed ESCs (**Figure 8 and S10B-C**). For *Sox2*, MLL2 secures a unique enhancer-promoter loop in both naïve and primed ESCs but while loss of *Mll2* is compensated by another loop in naïve ESCs, and so no significant change in chromatin mobility or in transcription, this is not the case in primed ESCs, where loss of *Mll2* disrupts the enhancer-promoter loop completely, leading to an increase in chromatin mobility and a decrease in transcription. The transcriptional amplitude of *Sox2* is associated with dynamic interactions between the gene and super-enhancer condensates (*90*) so MLL2’s ability to stabilise enhancer-promoter loops likely plays a key role, increasing the encounter frequency between the gene and its nearby super-enhancer condensates. However, these dynamic interactions are clearly different in naïve and primed ESCs for the *Sox2* gene, leading to the observed cell state-specific effects at this gene.

The finding that MLL2 is required for a developmental transition towards neuroectoderm, but not for ESC self-renewal, resonates with findings for the architectural organisation mediated by CTCF and cohesin, which also play a limited role in transcription during self-renewal (*91, 92*) but is required for cellular transitions and embryo development (*93*). Moreover, like MLL2, CTCF looping increases the transcriptional amplitude of *Sox2* (*90*). The MLLs may therefore participate alongside the CTCF/cohesin management of the 3D genome or may present an overlapping web that contributes to stabilising lineage commitment decisions.

Interestingly, Platania et al (2024) showed that the *Sox2* promoter exhibits more restrained chromatin mobility during transcription than would be expected from just CTCF/cohesin looping and that this additional restriction of chromatin mobility homogenises and increases the frequency of enhancer-promoter encounters during transcription (*84*). However, they did not identify additional molecular players that form looping interactions and restrain chromatin mobility at the *Sox2* locus. Since CTCF/cohesin loops constantly form and fall apart (*94*), we propose a model where MLL2 maintains loops to restrict chromatin mobility and drive higher enhancer-promoter encounter frequency in the absence of CTCF/cohesin (**Figure 8I**). This could explain why CTCF depletion has minimal effect on *Sox2* transcription in naïve ESCs (*84*). In our proposed model (**Figure 8I**), MLL2 loops may also stall loop extrusion which Platania et al (2024) propose as a mechanism for constraining chromatin mobility further. Consistent with this model, we observe stabilisation of CTCF/cohesin loops genome-wide in naïve ESCs upon loss of *Mll2* (**Figure 6B**). So why does loss of *Mll2* destabilise *Sox2*’s enhancer-promoter loop in primed ESCs (**Figure 8B**)? This may be related to observations by Pekowska et al (2018) that there is a loss of short-range and a gain of long-range CTCF/cohesin loops as ESCs differentiate (*95*). It is possible that, in primed ESCs, disappearance of the shorter CTCF/cohesin loops lead to MLL2 now being required to stabilise *Sox2*’s enhancer-promoter loop and therefore maintain *Sox2* expression as ESCs prime and then differentiate (**Figure 8I**). It would therefore be interesting to investigate the interplay between CTCF/cohesin-driven loop extrusion and MLL2-associated looping in future studies.

Our findings underscore the need to look beyond histone marks when studying enhancer-promoter communication, as histone marks can change independently of 3D genome organization. Moreover, they reveal a non-canonical role for MLL2 in stabilizing enhancer-promoter loops that warrants further exploration. Future studies could explore how MLL2’s chromatin binding domains and protein-protein interactions influence chromatin mobility and differentiation. MLL2 is also one of four MLLs (KMT2A-D), all of which play roles in cell differentiation. Because all four MLLs and the *Drosophila* orthologs Trithorax and Trr/Lpt have a similar protein architecture, we anticipate that the findings for MLL2 will be relevant for the other MLLs. In particular, MLL1, the closest paralogue of MLL2, is similarly required for lineage commitment in several somatic stem cells. In intestinal stem cells, MLL1 is required for maintenance and for controlling lineage specification of intestinal secretory Paneth and goblet cells (*26, 67, 68*). From this perspective, it is likely that the other MLLs also restrain chromatin mobility and collectively may substantially account for enhancer-promoter loop stabilisation during differentiation in many stem cells and lineages. Recent evidence from ESCs enhances the proposition for MLL co-operativity (*96*).

## Limitations

Apoptosis is increased about 2-fold in *Mll2−/−* ESCs (*33*). Consequently, we selected single-cell RNA-seq and quantitative immunofluorescence thresholds to remove smaller apoptotic cells and the SMLM experiments were designed to avoid fields of view containing apoptotic cells. However, it is possible that elevated apoptosis accounts for some of our observed changes. Our SMLM analysis was carried out over a 5-second timescale to monitor changes at the level of TADs and to avoid cell cycle- and transcription-dependent effects on chromatin mobility. It therefore remains possible that MLL2 also affects chromatin at different timescales. Alternatively, MLL2-dependent effects may be more pronounced at specific cell cycle phases since MLL2 acts primarily in G1 cells (*97*). We also cannot easily exclude transcription-dependent effects, although this seems unlikely because MLL2 affects chromatin mobility similarly in naïve and primed ESCs while having an opposing effect on transcription in these cell types. We also cannot exclude the possibility that MLL2 loss influences chromatin mobility indirectly. For example, MLL2-dependent loss of FOXD3 may lead to an increase in H3K27ac (chromatin decompaction), in SS18-BAF-induced chromatin remodelling or in recruitment of DNA methyltransferases (*36*).

## Materials and Methods

### Plasmids used

Plasmids used:
| Plasmid Name | Description | Source |
| --- | --- | --- |
| LZ10 PBREBAC-H2BHalo | Construct expressing the human H2b sequence with a C-terminal HaloTag under the HF1 promoter. Includes an ampicillin and a neomycin resistance cassette. | (98)<br>Addgene #91564 |
| pCyL43 (PBase) | Construct expressing PBase, the PiggyBac transposase. Includes an ampicillin resistance cassette. | (99) |
| PB-TRE3G-dCas9-eGFP-WPRE-ubcp-rtTA-IRES-puroR | Construct with dual promoter backbone, with a TRE3G (Tet-on) promoter expressing GFP-tagged inactive dCas9 and the ubiquitin C promoter expressing the reverse tetracycline-controlled transactivator, rtTA, and a puromycin | (82) |
|  | cassette via an IRES sequence. |  |

### Mouse embryonic stem cell (mESC) lines

**Mouse embryonic stem cell (mESC) lines:**
| Cell line | Description | Source |
| --- | --- | --- |
| *Mll2F/F;CreERT2/+* | ESCs established *de novo* from blastocysts with LoxP sites either side of exon 2 of the endogenous *MLL2* locus and expressing CreERT2 recombinase from the Rosa26 locus. MLL2 cKO induced by addition of 4-OHT (4-hydroxy-tamoxifen) for 48h and validated by genomic PCR and Western blot (**Figure S1A**) | (35) |
| *Mll2F/F;CreERT2/+ MLL2-GFP* | *Mll2F/F;CreERT2/+* ESCs described above with stably integrated BAC expressing GFP-tagged MLL2. Addition of 4-OHT leads to depletion of endogenous but not GFP-tagged MLL2. | (35) |
| *Mll2F/F;CreERT2/+ MLL2(N2650A)-GFP* | *Mll2F/F;CreERT2/+* ESCs described above with stably integrated BAC expressing GFP-tagged MLL2 where MLL2 contains a N→A point mutation at position 2650, resulting in loss of catalytic activity. Addition of 4-OHT leads to depletion of endogenous but not GFP-tagged MLL2 (N2650A). | This study |
| *Mll2F/F;CreERT2/+ H2b-HaloTag* | same as *Mll2F/F;CreERT2/+* ESCs described above but with an additional stably integrated piggyBac construct carrying *H2b* with a C-terminal HaloTag. MLL2 cKO again induced by addition of 4-OHT | This study |
| *Mll2F/F;CreERT2/+ MLL2(N2650A)-GFP* | *Mll2F/F;CreERT2/+ MLL2(N2650A)-GFP* ESCs described above but with an additional | This study |
| *H2b-HaloTag* | stably integrated piggyBac construct carrying *H2b* with a C-terminal HaloTag. |  |
| *Mll2F/F;CreERT2/+ dCas9-GFP* | *same as Mll2F/F;CreERT2/+ ESCs described above but with an additional stably integrated piggyBac construct carrying dCas9 with a C-terminal GFP tag.* | This study |

### Sequencing datasets

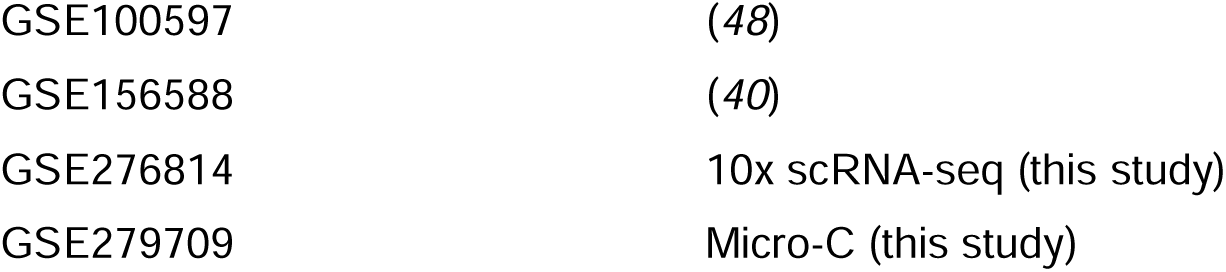

### Software

#### Data collection

|  |  |
| --- | --- |
| Microscope image acquisition | Micro-manager (100) |
|  | (https://www.micro-manager.org) |
| ImageJ software |  |
|  | (101, 102) |

#### Data analysis – sequencing

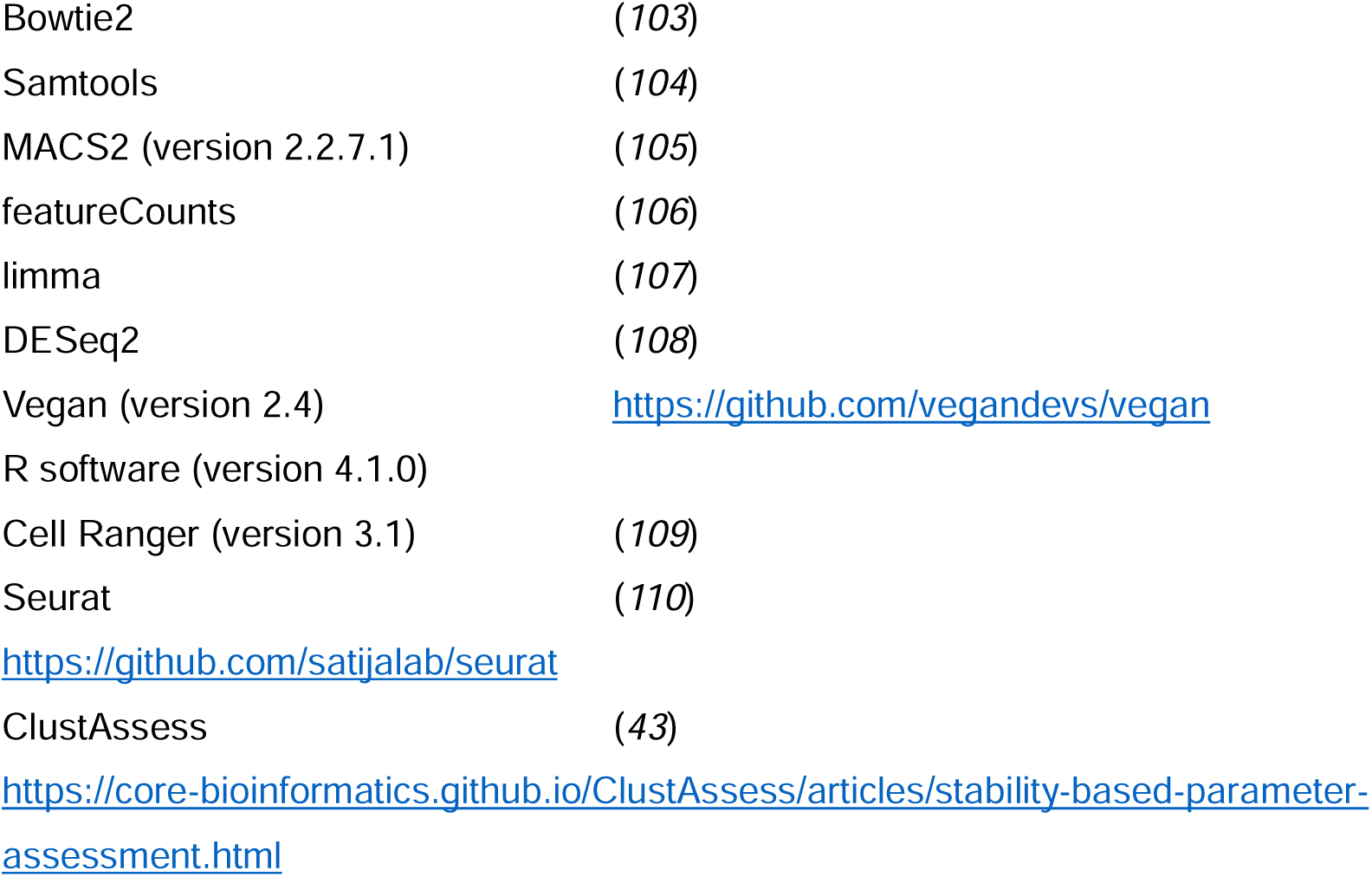
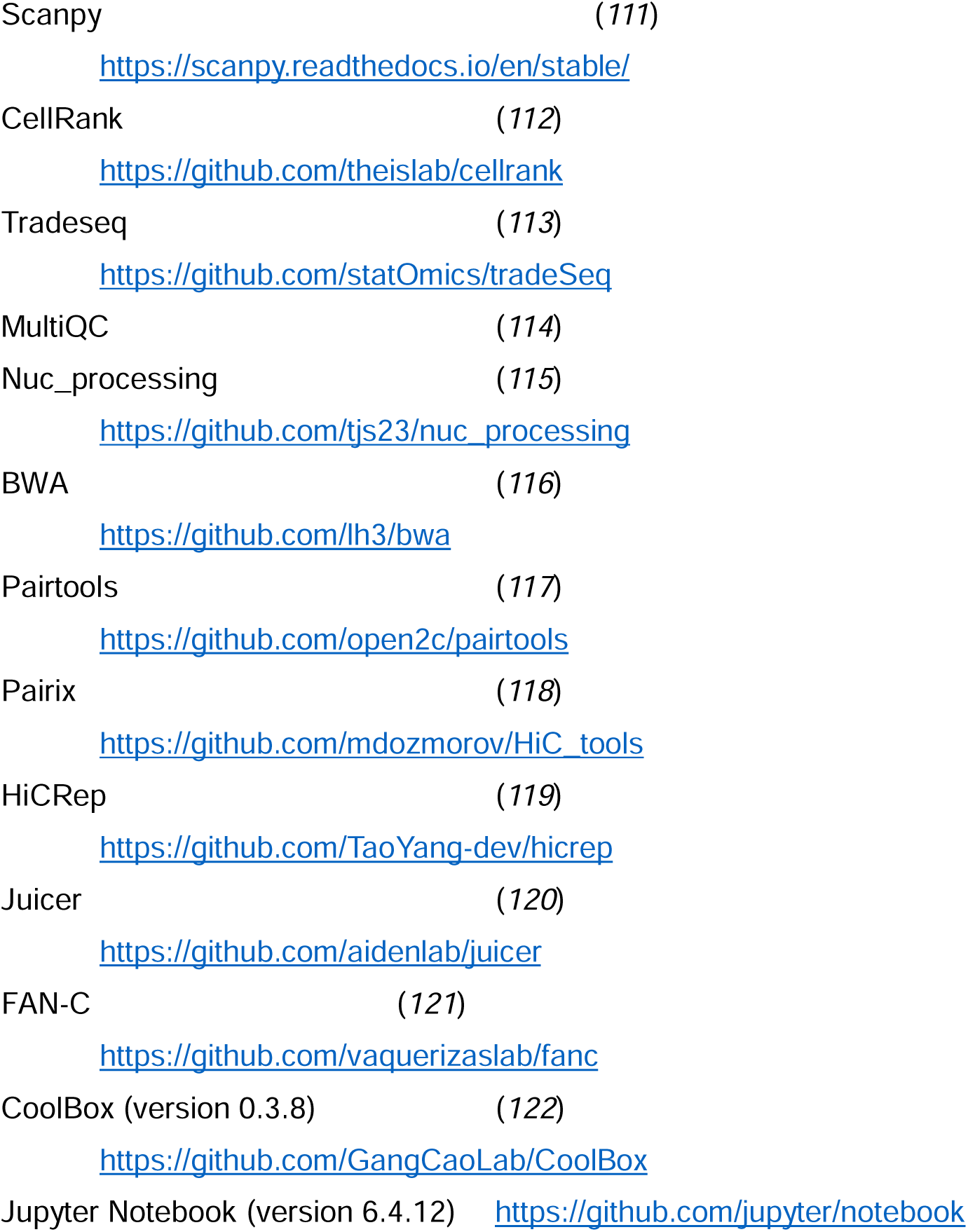

**Data analysis - sequencing**
|  |  |
| --- | --- |
| Bowtie2 | (103) |
| Samtools | (104) |
| MACS2 (version 2.2.7.1) | (105) |
| featureCounts | (106) |
| limma | (107) |
| DESeq2 | (108) |
| Vegan (version 2.4) | https://github.com/vegandevs/vegan |
| R software (version 4.1.0) |  |
| Cell Ranger (version 3.1) | (109) |
| Seurat | (110) |
|  | https://github.com/satijalab/seurat |
| ClustAssess | (43) |
|  | https://core-bioinformatics.github.io/ClustAssess/articles/stability-based-parameter-assessment.html |

|  |  |  |
| --- | --- | --- |
| Scanpy | (111) | https://scanpy.readthedocs.io/en/stable/ |
| CellRank | (112) | https://github.com/theislab/cellrank |
| Tradeseq | (113) | https://github.com/statOmics/tradeSeq |
| MultiQC | (114) |  |
| Nuc_processing | (115) | https://github.com/tjs23/nuc_processing |
| BWA | (116) | https://github.com/lh3/bwa |
| Pairtools | (117) | https://github.com/open2c/pairtools |
| Pairix | (118) | https://github.com/mdozmorov/HiC_tools |
| HiCRep | (119) | https://github.com/TaoYang-dev/hicrep |
| Juicer | (120) | https://github.com/aidenlab/juicer |
| FAN-C | (121) | https://github.com/vaquerizaslab/fanc |
| CoolBox (version 0.3.8) | (122) | https://github.com/GangCaoLab/CoolBox |
| Jupyter Notebook (version 6.4.12) |  | https://github.com/jupyter/notebook |

#### Data analysis – imaging

***Data analysis - imaging***
|  |  |  |
| --- | --- | --- |
| CellProfiler | (123) | https://cellprofiler.org/ |
| 3D SMLM peak fitting | (124) | https://sourceforge.net/projects/easy-dhpsf/ |
| Trajectory analysis | (125) | https://github.com/wb104/trajectory-analysis |
| SMLM Analysis Toolbox | (this study) | https://github.com/stanleystrawbridge/singleMoleculeLocalizationMicroscopyAnalysisToolBox |
| Trajectory classification | (71) | https://zenodo.org/record/6497411#.YmlGFy8w3q0 |

### 1 Cell culture

#### Embryonic stem cell (ESC) line generation and validation

MLL2 Cre-inducible conditional knockout (cKO) ESCs were generated as previously described (*35*). Tamoxifen was added for 2 days to generate knockout ESCs. Genomic PCR and Western blots ensured gene knockout and protein knock-down respectively. Cell numbers were doubled in knockout conditions to ensure similar numbers of cells when compared to wild-type conditions after 2 days.

#### Primary antibody for Western blot

Primary antibody for Western blot:
| Antigen | Dilution | Host | Isotype | Manufacturer |
| --- | --- | --- | --- | --- |
| MLL2 | 1:1000 | Rabbit | IgG1 | (32) |

#### Secondary antibody for Western blot

Secondary antibody for Western blot:
| Fluorophore | Dilution | Host species | Reactivity | Manufacturer | Cat # |
| --- | --- | --- | --- | --- | --- |
| HRP | 1:10000 | goat | anti-rabbit | Thermo Scientific | 31460 |

MLL2 cKO cells expressing GFP-tagged MLL2 or GFP-tagged MLL2 N2650A (catalytically-dead) ectopically were generated by first using recombineering to generate a bacterial artificial chromosome (BAC) containing the *Mll2* gene (Source Biosciences, bMQ-389B2) tagged with GFP internally inserted on the N-terminal part 5’ upstream of the Taspase cleavage sites (*126*). An internal ribosomal entry site (IRES) followed by a neomycin resistance gene was placed 3’ downstream of the open reading frame. The N2650A mutation was then introduced to this BAC by site-directed mutagenesis. 1 μg of purified BAC DNA was transfected into ESCs using Lipofectamine LTX and Plus reagent (Invitrogen, 15338-100). Cells were selected with 175 μg/ml G418 starting 1 day after lipofection and picked clones were validated by Western blot and microscopy. The GFP-tagged MLL2 N2650A cells were validated by histone chromatin immunoprecipitation (ChIP) followed by qPCR. 1.5×108 cells on five 15 cm dishes were washed with PBS and crosslinked with 10 ml/dish 1% formaldehyde/DMEM+Glutamax for 20 min at RT. The formaldehyde was quenched by adding 500 µl/dish of 2.5 M glycine and the crosslinked cells were washed with cold PBS, scraped from the dishes in cold PBS, collected by centrifugation with 600 xg for 5 min at 4°C and pellets frozen at 80°C until further processing. The ChIP was performed as previously described (*35*) with the following adaptation: the sonicated supernatant (input DNA) was equally divided up into 3 to 5 separate tubes and incubated with 7 µg of H3K4me1/me3– specific antibodies (Diagenode C15410037, Abcam ab8580). To normalize histone ChIPs against the background binding of histones to beads, a control without antibodies was always performed in parallel. Primers for the histone ChIP-qPCR were:

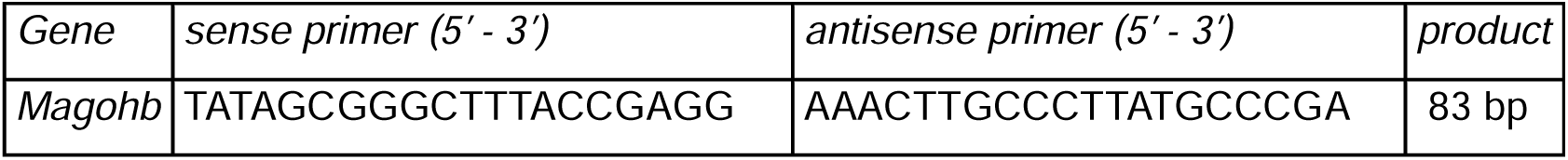

MLL2 cKO ESCs expressing HaloTag-tagged histone H2B (+/− GFP-tagged MLL2 N2650A) were generated by transfecting cells with the LZ10 PBREBAC-H2BHalo (Addgene, Plasmid #91564) (*98*) and PiggyBAC transposase PBase plasmids (*99*). 1 µg of the purified plasmids were added to freshly passaged MLL2 cKO ESCs using Lipofectamine-3000 (Thermo Fisher, #L3000001) following the standard protocol. G418-resistant mESCs were selected by growing cells in the presence of 1 μg/ml G418. Single colonies were picked after 7 days, plated individually, and grown to confluence. Clones were chosen for experiments if they had positive nuclear signal after labelling with JFX_549_ or with PA-JF_549_ and if immunofluorescence experiments confirmed that they proceeded through priming as expected.

MLL2 cKO ESCs expressing GFP-tagged inactive dCas9 were generated by transfecting cells with the PiggyBAC transposase PBase plasmid (*99*) and the PB-TRE3G-dCas9-eGFP-WPRE-ubcp-rtTA-IRES-puroR vector containing a dual promoter backbone, with a TRE3G (Tet-on) promoter expressing GFP-tagged inactive dCas9 and the ubiquitin C promoter expressing the reverse tetracycline-controlled transactivator, rtTA, and a puromycin cassette via an IRES sequence (*82*). Puromycin-resistant ESCs were selected for 7 days and doxycycline added for 24 hours to induce expression of dCas9-GFP (through activation of the rtTA). Stable transfectants were FACS sorted for GFP expression.

#### ESC culture conditions

Naïve ESCs were grown in 2iLIF medium at 37 °C, 7% CO_2_ and 80% humidity as previously described (*10*). Phenol red-free 2iLIF medium was prepared fresh every week by adding 1 µM PD0325901 (ABCR, AB 253775), 3 µM CHIR99021 (ABCR, AB 253776) and 10 ng/ml murine leukaemia inhibitory factor (mLIF, provided by the Department of Biochemistry, University of Cambridge) to N2B27 basal medium. N2B27 medium is composed of 50% phenol red-free DMEM/F-12 (Gibco #21041025) and 50% phenol red-free and L-Glutamine-free Neurobasal (Gibco #12348017) supplemented with 0.5x N2 (made and batch tested in-house by Cambridge Stem Cell Institute), 1x B-27™ Supplement (Gibco #17504044), 2mM L-Glutamine (Life tech, #25030024), and 0.1 mM 2-mercaptoethanol (in-house/Life Tech, #1985023). Cells were passaged every two days by washing in PBS (Sigma-Aldrich #D8537), followed by incubation with accutase (Sigma-Aldrich #A6964) for 2 min at 37 C to detach. Cell clumps in suspension were then washed and pelleted in PBS before resuspending and re-plating as single-cell suspension in fresh medium. To help cells attach to the surface, plates were incubated for 15 minutes at 37 C in PBS containing 0.2% gelatin (Sigma-Aldrich #G1890). All cell lines were routinely screened for mycoplasma contamination at least twice yearly and tested negative.

#### Priming of naïve ESCs to primed epiblast-like cells (EpiLCs)

Naïve ESCs were converted to primed EpiLCs as previously described. Naïve mESCs were converted to primed EpiLCs as previously described (*14*). Briefly, plates were coated with 20 µg/ml human plasma fibronectin (Sigma-Aldrich #FC010-10MG) for 5 min, then incubated to dry for 30 min at 37°C and 7% CO_2_ prior to plating cells at a density of 3×10^5^ cells per 6-well (∼3×10^4^ cells/cm^2^). 2i inhibitors and mLIF were removed and replaced with N2B27 basal medium supplemented with 12.5 ng/ml FGF2 and 20 ng/ml Activin A (both provided by the Department of Biochemistry, University of Cambridge). Cells were cultured in FGF2 plus Activin A medium for three days with daily media changes.

#### Differentiation of EpiLCs to neuroectoderm

To continue differentiating EpiLCs to neuroectoderm, wells were washed with DMEM/F12 and cells were cultured on coverslips for 7 days in N2B27 without growth factors. The medium was changed every other day. After 7 days in N2B27, coverslips were fixed and permeabilized for immunofluorescence. Neural rosettes from 6-well plates were dissociated with 0.5 ml accutase for 5 min at 37°C, diluted and resuspended with 4.5 ml DMEM/F12, collected by centrifugation for 5 min at 215 g and cultured on non-adhesive 6 cm suspension culture dishes in 4 ml NSA with 0.5x B27 for 2-4 days to allow formation of neurospheres. These were then transferred to centrifugation tubes, allowed to settle down for 5 min and the old medium was carefully aspirated. Spheres were resuspended in 3-4 ml fresh NSA with B27 and seeded to 6-well plates (Nunc) or 6 cm dishes (VWR) that were beforehand coated with 0.1% gelatin for at least 1 hr at RT, aspirated and dried under the laminar flow hood with open lids for 10-20 min. Spheres were allowed to attach and grow out for 2 days without moving the dishes around before still unattached spheres were aspirated and medium was changed. When reaching near confluency (95%) at day 3-5 on gelatine, neural stem (NS) cells were passaged for the first time by aspirating the medium and detaching the cells with 0.5 ml Accutase for 5 min at 37°C. Accutase was diluted with 4.5 ml DMEM/F12, cells were collected by centrifugation for 5 min at 215 g,resuspended in fresh NSA with B27 and seeded to gelatin-coated dishes (Nunc or WVR) in a splitting ratio of 1:3 to 1:8. B27 was removed from the medium one day after the first passage and NS cells were regularly passaged every 3-5 days shortly before confluency was reached (95%) as described above. EpiSCs were differentiated similarly but after 21-35 days of culture in FGF2 plus Activin A medium, passaging cells every 3-5 days around 24 h after reaching full confluency.

### 2 Assays

#### Immunofluorescence

##### Immunofluorescence staining of neural rosettes

Cells on glass cover slips were washed with PBS, fixed 10 min in 4% formaldehyde/PBS (Merck Millipore, 104003), washed with PBS, permeabilized 15 min in 0.5% Triton X-100/PBS (Sigma-Aldrich, T8787), washed again with PBS and blocked 30 min with 3% BSA/PBS (Sigma-Aldrich, A7906). Cover slips were either stored in 3% BSA/PBS (Sigma-Aldrich, A7906) for up to 3 months at 4 °C or processed directly for immunofluorescence analysis as follows. Each cover slip was incubated in 25 μl drops of primary antibodies diluted in 10% goat serum/PBS (Sigma-Aldrich, G9023) over night at 4 °C in a humidified atmosphere. The following day cover slips were washed in PBS and incubated 1-4 hours in 25 μl drops of appropriate secondary antibodies diluted in 10% goat serum/PBS (Sigma-Aldrich, G9023) at 4°C in the dark. Cover slips were washed again in PBS and stained with 100 ng/ml 4,6-diamidino-2-phenylindole (DAPI)/PBS for 2 min (Sigma-Aldrich, D8417). After washing in PBS and then in ddH2O each cover slip was mounted to glass slides with 7 μl Mowiol (Sigma-Aldrich, 81381) and dried overnight at RT before imaging with the TCS SP5 confocal system.

##### Immunofluorescence staining of priming transition

Cells were cultured in Fibronectin-coated Ibidi 8-well micro-chambers (Ibidi, IB-80826). Cells were washed in PBS and fixed with 4 % PFA for 10 mins at room temperature. Fixed cells were rinsed three times in PBS and then blocked for two hours at room temperature in blocking buffer: PBS containing 0.1 % TritonX-100 (Sigma-Aldrich, T8787-100M) and 3% normal Donkey Serum (Sigma-Aldrich, D9663-10ML). Cells were incubated with primary antibodies diluted in blocking buffer (see concentrations below) at 4 °C overnight, followed by three 10 min washes in PBST (PBS containing 0.1% TritonX-100). Incubation with secondary antibodies and 1000 ng/ml DAPI (Sigma-Aldrich, MBD0015-1ML) diluted in blocking buffer (see concentrations below) was performed for two hours at room temperature in the dark. Stained samples were then rinsed in PBST three times for 10 min before being stored in PBS at 4 °C overnight prior to imaging on a Nikon Spinning Disk Ti inverted microscope using a 40x oil immersion objective and an Andor iXon + camera.

###### Primary Antibodies for immunofluorescence

| Antigen | Dilution | Host | Isotype | Manufacturer | Cat # | LOT # |
| --- | --- | --- | --- | --- | --- | --- |
| OTX2 | 1:50 | goat | IgG polyclonal | R&D Systems | AF1979 | KNO0918121 |
| NANOG | 1:200 | rat | IgG2a, kappa | eBioscience | 14-5761-80 | 2003079, 2403803, 2269925 |
| SOX2 | 1:200 | rat | IgG2a | eBioscience | 14-9811-82 | 2213316 |
| PAX6 | 1:20 | mouse | IgG1 | DSHB | supernatant |  |
| NESTIN | 1:20 | rat | IgG1 | DSHB | Rat-401 |  |
| ZO-1 | 1:100 | rabbit | IgG1 | BD Biosciences | 610966 |  |

###### Secondary Antibodies for immunofluorescence

Secondary Antibodies for immunofluorescence:
| Fluorophore | Dilution | Host species | Reactivity | Manufacturer | Cat # | LOT |
| --- | --- | --- | --- | --- | --- | --- |
| DyLight488 | 1:500 | donkey | anti-goat | DyLight | SA5-10086 | WE3275103 |
| AF555 | 1:500 | donkey | anti-rat | Abcam | ab150154 | GR3262493-3 |
| CF647 | 1:1000 | donkey | anti-rabbit | Biotium | 20047-1mg | 17C0123 |

###### Microscope Settings

Microscope Settings:
| Fluorophore | Excitation Wavelength | Laser Power | Exposure Time |
| --- | --- | --- | --- |
| DAPI | 405 nm | 50.0% | 80 ms |
| DyLight488 | 488 nm | 20.0% | 200 ms |
| AF555 | 561 nm | 20.0% | 400 ms |
| CF647 | 638 nm | 68.1% | 300 ms |

#### Single-cell Transcriptomics (10X Genomics)

##### Cell Harvesting

MLL2 cKO and control cells were transitioned from naive to primed pluripotency as described above. Cells harvested at different time points during the protocol (0 (2iLIF), 24, 48 or 72 hours) were prepared for 10X genomic sequencing, according to the manufacturer’s protocol. The experiment was repeated twice on separate days. Briefly, cells were washed twice with PBS, detached and collected with PBS supplemented with 0.4 mg/mL BSA (Sigma-Aldrich, A2153) to neutralise excess accutase activity. After pelleting and resuspension in PBS/BSA, live cells were counted using a TC20^TM^ automated cell counter (BioRad Cat.# 1450102), BioRad Cell Counting Slides (BioRad #1450011) and 0.4% Trypan Blue Solution (Gibco^TM^ #15250061). Cells were washed twice with PBS/BSA and passed through a 40 µm cell strainer (Greiner, 542040), then counted, pelleted and resuspended in PBS/BSA at a final concentration of 40 cells/µl. After harvesting, cells were kept on ice between handling and all pipetting steps except the final resuspension step were carried out using wide-bore pipette tips to minimise shearing forces/cell damage.

##### Single-cell RNA-seq library preparation and sequencing

Libraries were prepared and sequenced in the Cancer Research UK Cambridge Institute Genomics Core Facility using Chromium Next GEM Single Cell 3′ GEM, Library & Gel Bead Kit v3.1 (10X Genomics, #PN-1000121), Chromium Next GEM Chip G Single Cell Kit (10X Genomics, #PN-1000120) and Chromium Next GEM Single Cell 3 Reagent Kits User Guide v3.1 (Manual Section CG000204 Rev D; 10X Genomics). Cell suspensions were loaded on the Chromium instrument with the expectation of collecting gel-beads emulsions containing single cells. RNA from the barcoded cells for each sample was subsequently reverse-transcribed in a C1000 Touch Thermal cycler (Bio-Rad) and all subsequent steps to generate single-cell libraries were performed according to the manufacturer’s protocol with no modifications. cDNA quality and quantity were measured with Agilent TapeStation 4200 (High Sensitivity D5000 ScreenTape, Reagents and Ladder, Agilent #5067-5592, −5593 and −5594) after which 25% of material was used for gene expression library preparation. Library size distribution was confirmed with Agilent TapeStation 4200 (High Sensitivity D1000 ScreenTape, Reagents and Ladder, Agilent #5067-5584, −5585, −5587 and −5603) and Qubit 4.0 Fluorometer (ThermoFisher Qubit™, dsDNA HS Assay Kit, #Q32853) to quantify dsDNA. Each sample was normalised and pooled in equal molar concentration. To confirm concentration, pools were qPCRed using the KAPA Library Quantification Kit (Roche # KK4824) on a QuantStudio 6 Flex Real-Time PCR System (Applied Biosystems, #4485691) before sequencing. Pools were sequenced on an Illumina NovaSeq6000 sequencer with the following parameters: 28 bp, read 1; 8 bp, i7 index; and 91 bp, read 2.

#### Micro-C

Micro-C was performed using the Dovetail Micro-C Kit (Dovetail Genomics, #21006) following the manufacturer’s instructions with slight modifications, as described below. The analysis consisted of two independent experiments, SLX-22775 and SLX-23477/SLX-23479, each with technical duplicates, and with the latter including the catalytic activity mutant (referred to as N2650A).

##### Cell Harvesting

Briefly, 1 million naïve or primed pluripotent cells were harvested, washed in 1x PBS, pelleted and the pellets placed at −80°C for at least 30 min before proceeding with the protocol.

##### Crosslinking and MNase digestion

Long-range chromatin crosslinking was achieved by addition of DSG (disuccinimidyl glutarate; ThermoFisher Scientific #A35392), followed by 37% formaldehyde (Sigma-Aldrich, #F8775) to stabilise shorter-range DNA-protein and protein-protein interactions. After two wash steps, cells were subjected to *in situ* micrococcal nuclease (MNase) digestion. We used 2 uL of MNase enzyme in all experiments. EGTA (ethylene glycol-bis(β-aminoethyl ether)-N,N,N′,N′-tetraacetic acid) was added to terminate the reaction after exactly 15 min. Cells were subsequently lysed using SDS (sodium dodecyl sulphate) and the lysate stored at −80°C until the next step.

##### Quality control of MNase digestion profile (QC 1)

For the first quality control step, we assessed the size distribution of the MNase-digested DNA to ensure it was the recommended 40-70%. A small sample taken from the lysate was subjected to crosslink reversal and Proteinase K digestion to remove protein contaminants. DNA was purified using the Zymo Research DNA Clean and Concentrator-5 Kit (Zymo Research, #D4013) and analysed using an Agilent D5000 ScreenTape (Agilent, #5067-5588) on the Agilent TapeStation 4200 (JCBC NGS facility; Analysis Software Version 5.1). DNA concentration was determined using a Qubit 2.0 Fluorometer (JCBC NGS facility; Qubit^TM^, ThermoFisher Scientific).

##### Proximity ligation, end polishing and crosslink reversal

To carry out proximity ligation, samples were thawed, and 1000 ng of each lysate immobilised on chromatin capture beads. Following two washes, an end polishing step was performed to fill ‘sticky ends’ generated by MNase digestion. After another wash, crosslinked fragments were ligated in a two-step reaction using the provided bridge, bridge ligase and the intra-aggregate ligation reagents. Consequently, a combined crosslink reversal and Proteinase K digestion step was carried out and resulting ligated fragments were eluted from the beads. This step also introduced a biotin-label into the ligated fragments, crucial for enrichment of ligated fragments using streptavidin beads, as described below. DNA was subsequently purified using SPRIselect beads (Beckman Coulter # B23319) at a 1.8-fold bead-to-sample ratio to ensure maximum recovery of all DNA fragments ≥100 bp and stored overnight at −20 °C.

##### End repair, USER digest and library indexing

Purified and proximity-ligated fragments were subjected to End Repair, followed by Ligation with Illumina Adapters, USER (uracil-specific excision reagent) digest and another SPRIselect bead-based purification (2.2 bead-to-sample ratio). Ligated sample-adapter fragments were then immobilised on streptavidin beads (via the biotinylated bridge introduced before) and washed for the indicated times with several provided buffers (LWB, NWB, 1x Wash Buffer). Samples were indexed using a set of eight unique UDI Primer Pairs in a standard PCR reaction (12 cycles) as recommended. Primer Pairs were allocated as follows:

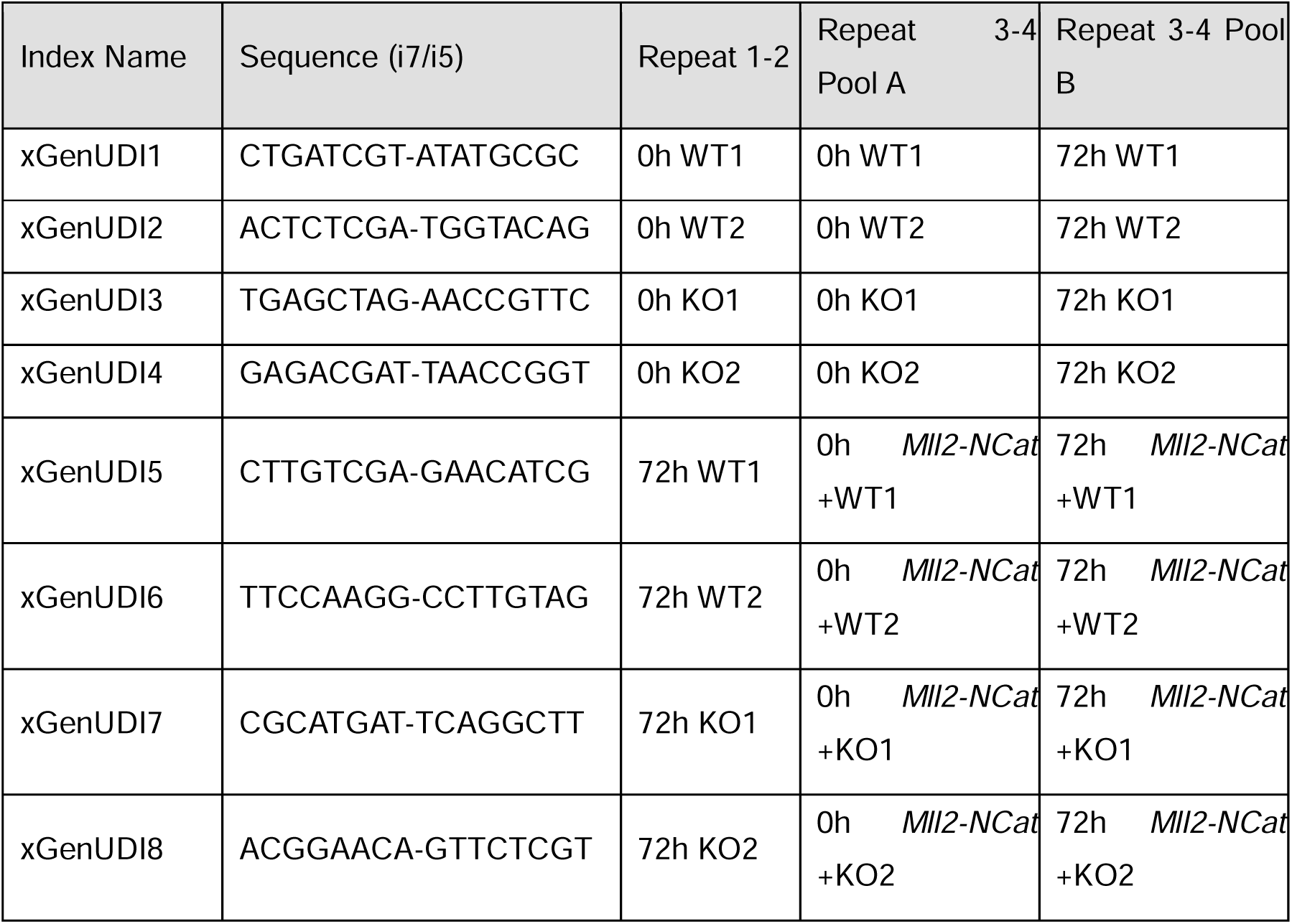

The resulting libraries were subjected to a double-sided selection using SPRIselect beads (right-sided selection at 0.5x, followed by left-sided selection of supernatant at 0.7x bead-to-sample ratio), which excludes any fragments sized <350 bp or >1000 bp.

##### Quality control for library size distribution (QC2)

A small sample of each library was run on an Agilent TapeStation 4200 to validate that we had the required fragment size distribution and a Qubit 2.0 Fluorometer to determine DNA concentration.

##### Library pooling

Eight libraries (corresponding to UDI primer pairs 1-8) were pooled at equal concentration to yield a final concentration of 5 nM and the average fragment size of the pooled library was determined using the Agilent TapeStation 4200. Average pooled library lengths are listed below:

| Library Name | Library Barcode | Samples | Average Length [bp] |
| --- | --- | --- | --- |
| A | SLX-23477 | 0h WT, 0h KO, 0h *MII2-NCat*+WT, 0h *MII2-NCat* +KO | 460 |
| B | SLX-23479 | 72h WT, 72h KO, 72h *MII2-NCat* +WT,<br>72h *MII2-NCat* +KO | 468 |

##### Submission for sequencing

Samples were submitted for 200 bp paired-end sequencing at the CRUK-CI Genomics Core Sequencing Facility and sequenced on a NovaSeq S4 Flow Cell using a standard Illumina sequencing workflow.

#### Live-cell Single Molecule Localisation Microscopy (SMLM)

##### Cell preparation

Prior to imaging, cells were passaged onto fibronectin-coated (16.7 μg/ml; Sigma-Aldrich, #FC010; later #F0895) 35 mm glass bottom dishes No 1.0 (MatTek Corporation, #P35G-1.0-14-C) in phenol red-free 2i/LIF or FA medium, respectively, and grown for the indicated duration, according to the protocols described above.

##### Labelling of H2B-Halo expressing cells

On the day of imaging, H2B-Halo expressing ESCs were incubated with 50 nM PA-JF_549_ Halo ligand dyes (*74*) (dyes were a kind gift from Luke Lavis’s Lab, Janelia Research Campus; reconstituted in DMSO) for 15 min at 37 °C, followed by three PBS washes and incubation for 30 min at 37 °C to release excess dye, and then imaged in fresh medium. Fixed cells were prepared by washing cells with PBS, then incubating for 10 min in PBS containing 4% formaldehyde (Thermo Fisher Scientific, 28908) and then washing in PBS three times.

##### Labelling of *Sox2* gene in dCas9-GFP expressing cells

To label the *Sox2* gene, we first induced expression of dCas9-GFP by adding 1 µg/ml doxycycline a day before cells were plated onto imaging dishes containing 2i/LIF or FA medium. Cells were then transfected using lipofectamine CRISPRMAX™ Cas9 Transfection Reagent (Thermo Fisher Scientific, CMAX00008) with Atto647-labelled gRNAs as cells were plated onto imaging dishes. We targeted the dCas9 to a genomic repeat (15 repeats) that is 1 kb upstream of the *Sox2* gene promoter using 20 µM of a custom crRNA (GGGAGGGAGGAAGUGGGAGG; Sigma, TRACRRNAMOD) assembled with Alt-R™ CRISPR-Cas9 tracrRNA, ATTO™ 647 (IDT, 10007853). crRNA and trcRNA were annealed for 5 min at 95 °C, then on ice for 20 min, prior to transfection.

##### Microscope setup

A custom-built double helix point spread function (DHPSF) microscope was used for 3D single-molecule tracking as previously described (*75, 127*). Briefly, lasers were used at appropriate wavelengths for fluorophore activation (405 nm: 100 mW, 06-MLD, Cobolt) and excitation (561 nm: 200 mW, 06-DPL, Cobolt) of the PA-JF_549_ fluorophore. Atto647-gRNAs were excited using a 638 nm laser (180 mW, 06-MLD, Cobolt). Laser beams were circularly polarised, expanded, collimated and focused to the back focal plane of an index-matched water immersion objective lens (CFI Plan Apo IR 60XC WI 1.27 NA, MRD07650, Nikon) mounted on an inverted fluorescence microscope (Eclipse Ti2, Nikon).

The microscope was equipped with a built-in Perfect Focus System, providing automated focusing over extended imaging periods and precise axial movement. This allows for accurate data acquisition and calibration of the DHPSF microscope by adjusting the axial position of the objective lens. A high-precision motorised linear XY stage (HLD117NN, Prior) was used to position the imaging sample. The system was also equipped with an incubator chamber for live-cell imaging (Digital Pixel), enclosing the stage and objective in a sealed plastic chamber with two heaters to consistently maintain the temperature at 37°C.

##### Sample illumination

At the beginning of each day of experiments, the laser beam was aligned to the Highly Inclined and Laminated Optical sheet (HILO) illumination mode for better optical sectioning, resulting in reduced fluorescence background, improved signal-to-noise ratio (SNR), and reduced photobleaching of the sample. The thickness of the HILO illumination slice was similar to that of the DH axial range (∼4 µm). Once optimal HILO alignment was achieved, constant illumination was maintained throughout all experimental conditions.

##### Sample emission

The emission signal was redirected from the excitation path into the emission path by a quad-band dichroic mirror (Di01-R405/488/561/635-25×36, Semrock). The detection arm included a 4f system with two 200-mm tube lenses (TTL200-A, Thorlabs) and a 580nm or 687 nm double-helix phase mask (Double Helix Optics) positioned at the Fourier plane to generate the axially-modified PSF containing 3D information. The resulting signals were then focused by a third tube lens (TTL200-A, Thorlabs) onto an EMCCD detector (Evolve Delta 512, Photometrics). Long-pass and band-pass filters (BLP02-561R-25 and FF01-580/14-25 for PA-JF_549_ and FF01-676/29-25 for Atto647, Semrock) were placed immediately before the camera to selectively transmit the emitted fluorescence while blocking unwanted wavelengths of light.

##### Data acquisition

Micro-Manager (version 1.4) was used to control the shutters, sample stage, camera, and imaging parameters during data acquisition (*100*). Calibration series for each DHPSF channel were acquired using TetraSpeck 0.1 μm Microspheres (Thermo Fisher Scientific, #T7279) via a custom Micro-Manager script (10 frames per 40 nm axial increment spanning a total axial range of 4 µm) before any imaging experiments. For each experiment, 2000 frames of fluorescence images were collected with an exposure time of 500 ms. Although the 405 nm laser was used to validate cells, it was not required for the experiment itself because spontaneous activation of the PA-JF_549_ fluorophore provided sufficient density of single H2B molecules. 561 nm and 638 nm lasers were set to power densities of 40 and 0.7 W/cm^2^ respectively. Each experiment included at least 6 biological replicates, with each replicate covering 5 fields of view containing approximately 3 cells (total of 90 cells per condition).

### Statistical analysis

#### Analysis of MLL2-dependent cell fractions and genes using 10x scRNAseq

Raw sequencing reads were aligned to the mouse reference genome (mm10, 10x reference: refdata-gex-mm10-2020-A). Cell Ranger version 3.1 was used to generate unique molecular identified (UMI) count matrices for single cells using default settings (*109*). Seurat R package version 4.3.0 on R version 4.2.1 was used for data analysis for all conditions and both biological replicates (*110*). The UMI counts were normalised and scaled for each cell using a negative binomial regression-based heterogeneity removal method, *sctransform*. A principal component analysis (PCA) was performed on the normalised and scaled data. The Harmony algorithm was then used on the top 50 principal components, to integrate data from two biological replicates and reduce batch effects. After the integration step, cells were separated into clusters using Seurat’s shared nearest neighbour method, *FindNeighbors* and *FindClusters*, with resolution = 0.3. For visualisation and data exploration of cell-to-cell clustering and variation, the top 50 principal components with overlayed cluster information were plotted using a Uniform Manifold Approximation and Projection (UMAP) plot. The fraction of cells/cluster in wild-type and MLL2 cKO cells were compared using Fisher’s exact test.

##### Quality control

Parameter selection for the Seurat pipeline was optimised with the *ClustAssess* package version 0.3.1 (*43*). The *plot_feature_stability_boxplot* and *get_nn_importance* functions were used to assess UMAP clustering stability and consistency. 10 computational iterations (*n_repetitions*) were carried out with 50 principal components (*npcs*) and 50 nearest neighbours (*n_neighbours*).

<u>Differential gene expression</u> per cluster was performed using a negative binomial regression model. To identify MLL2-dependent genes per cluster, we used standard Seurat functions to conduct the analysis, first applying *PrepSCTFindMarkers* function to correct counts, then *FoldChange* to obtain log2-fold change for all genes in each cluster and *FindMarkers* to identify those that were differentially expressed. Differentially expressed genes (DEGs) were selected if they showed a p-value < 0.001. Since transcript levels/cell were not normally distributed, transcript levels were compared between conditions using Wilcoxon rank sum tests with Bonferroni correction. To assess whether MLL2-dependent genes had a likelihood of MLL2 binding, we overlapped our genes with a published MLL2 ChIP-seq (*35*) and then used a chi-squared test to assess whether there was an increase in the proportion of genes bound by MLL2 at DEGs versus at genes with no transcriptional change. We also assessed whether there were more MLL2 binding sites at DEGs versus at genes with no transcriptional change (within 1 kb of their gene regions) using a Mann Whitney test.

#### Comparison of scRNAseq with published scRNAseq datasets for validation

To compare our data to published single-cell datasets taken from mouse embryos *in vivo* and formative-like ESCs *in vitro*, we extracted the QC-processed, single cell count matrix data from GSE100597 (*48*) and GSE156588 (*40*) respectively. For each comparison, a list of common genes expressed in both datasets was identified and genes not expressed in both datasets were removed. For the *in vivo* comparison, we identified a list of the 3000 most-variable genes in the *in vivo* dataset using the *SCTransform* function from Seurat. For the *in vitro* comparison, we used the 3000 most-variable genes identified in our dataset (see above). We then re-applied our Seurat pipeline using the same functions, parameters and statistical tests as described above.

#### Trajectory analysis using Cell Rank

To visualize the time evolution of our system, we used the Python module *scanpy* (*111*), where we recomputed neighbours and UMAP coordinates with default parameters, excluding cells from the cluster of primarily apoptotic cells found at all timepoints. As a result, we obtained a continuous landscape. We then ran *cellrank* to produce a directional stream plot (*112*). Cellrank was initialised with the diffusion pseudotime kernel, also computed in *scanpy* (*128*). To test for differences in gene dynamics among MLL2 cKO and control cell conditions, we used the R package *tradeSeq* (*113*). Upon running the function *conditionTest*, we identified genes that have overall differential trajectories. Finally, we fed our 200 top differentially expressed genes to the function *clusterExpressionPatterns* and identified 15 clusters of genes that have similar patterns for both MLL2 cKO and control cell conditions.

#### Quantitative immunofluorescence data analysis

Data inspection and processing were performed in Fiji (is just ImageJ, version 2.9.0/1.53t; Java 1.8.0_322 [64-bit]) and involved adjustment of pixel display range (identical settings applied across conditions for each marker, except DAPI, which was auto-adjusted for best signal), cropping, conversion to RGB format and addition of scalebars.

For single-nucleus segmentation and quantification of signal, images were loaded into CellProfiler (version 4.2.5) and single nuclei were identified by DAPI signal (size range: 70-150 pixels; thresholding method: Otsu, thresholding strategy: global, lower and upper thresholding bounds: 0-1.0, thresholding smoothing scale: 1.3488). Binarised segmentation files were inspected manually and any erroneous conditions were excluded from further analysis. Average signal intensity was then measured by object, normalised by nuclear area and plotted using ggplot2 in RStudio (version 2022.02.3, R version 4.1.1). For calculation of fold-change values, we calculated the mean fluorescence per cell for each field of view. Since the mean fluorescence was normally distributed (∼10-100 cells/image), we compared means using an unpaired two-sided t-test. Technical background intensity for each channel was determined in a DAPI-only sample and processed in the same manner.

| Fluorophore | Mean background intensity | SD background intensity | n cells |
| --- | --- | --- | --- |
| DyLight488 | 0.001437 | 8.212416e-6 | 1982 |
| AF555 | 0.0007859 | 1.054071e-5 | 1982 |
| CF647 | 0.003491 | 1.710086e-4 | 1982 |

Mean background fluorescence intensity (normally distributed) was subtracted from mean fluorescence intensity per cell for each condition before calculation of fold-change.

#### ChIP-seq data analysis

ChIP-seq datasets of histone modifications (H3K4me3, H3K4me1, H3K27me3 and H3K27ac) at 0 and 72 hours during the priming transition were retrieved from the Gene Expression Omnibus (GEO) using the accession number GSE117896 (*16*). Raw reads were trimmed using *cutadapt* to remove adapters and low-quality regions with Phred score less than 20. Processed reads were then aligned against the mouse mm10 primary genome assembly using *bowtie2* with additional parameters *–end-to-end –very-sensitive* (*103*). Following the alignment, sam files were converted to bam format and sorted, duplicate-marked using *picard*, and filtered using *samtools view -F 2308 -q 30* to remove unmapped and low-quality alignment (*104*). Reads overlapping ENCODE problematic genomic regions were removed using *bedtools* (*129*). Regions with enriched signals (peaks) were identified for the blacklist-removed files using *macs3* (*105*). For H3K4me3 and H3K27ac marks, default parameters were applied for peak calling. For H3K4me1 and H3K27me3 marks, additional parameters --*broad --broad-cutoff 0.1* were used. Aligned reads and histone peaks were quality checked using the R package *ChipQC* (*130*). Aligned reads were extended to fragment lengths, and tracks of histone signals were generated using *deepTools bamCoverage --normalizeUsing CPM* (*131*). Lastly, the histone peaks at each time point were compared using *bedtools intersect* command (*129*) and filtered with respect to 2000 bp flanking window from Gencode annotated transcription start sites (TSS) (*132*). This enabled the categorisation of TSS-associated active promoters (H3K4me3 and H3K27ac) and bivalent promoters (H3K4me3 and H3K27me3), and active enhancers (H3K4me1 and H3K27ac).

ChIP-seq datasets of MLL2 binding (at 0 and 72 hours during the priming transition) were retrieved using the accession number GSE52071 (*35*) and processed using similar methods as histone datasets. In brief, trimmed reads were aligned using *bowtie2* and filtered to remove reads that were unmapped, low-quality, duplicates, or intersecting the ENCODE blacklisted regions (*103*). Peak calling was performed using *macs3* with default parameters (*105*), and signal tracks were produced using *deeptools*.

#### Micro-C data pre-processing

Quality control of raw files was assessed using *fastqc* (*114*). Barcode demultiplexing was performed using *nuc_processing* (*115*). Data preprocessing was performed following the Dovetail genomics pipeline (https://micro-c.readthedocs.io/en/latest/). In brief, raw reads were trimmed using *cutadapt* (*133*), and aligned to the mm10 mouse reference genome using *bwa* in MEM mode (*116*). *pairtools* (*117*) and *pairix* (*118*) were used to record valid ligation events and to index the pairs of genomic coordinates respectively to generate pair files.

#### Matrix binning and conformation analysis

For A/B compartment analysis and topologically associated domain (TAD) analysis, pairs files derived from the dovetail pipeline were binned into contact matrices and balanced with Knight-Ruiz (KR) method in *Juicer* (*120*). A/B compartments and TAD boundaries were then called using *FAN-C* (0.9.17) (*121*), applying default settings. HiCRep highlighted an acceptable level of consistency between replicates, and so replicates were merged (*119*). A/B compartment sizes, TAD sizes and TAD insulation scores were not normally distributed and therefore compared using a Mann-Whitney-Wilcoxon two-sided test with Benjamini-Hochberg correction.

Reproducibility of replicates was assessed using HiCRep (*119*), which highlighted an acceptable level of consistency for each cell type. Therefore, to increase the statistical power and detect chromatin interactions at higher resolution, replicates were pooled for the identification of statistically significant interactions. Pair files were combined and binned at 10 kb resolution using *cooler*. To account for differences in read depths, cool files were downsampled with respect to the dataset with the fewest intrachromosomal interactions. Loop calling was performed using FitHiC2 (v2.0.8) (*64*), with KR bias derived from HiCKRy.py command and parameters -U 2000000 --L 20000 to constrain extreme interaction sizes. To focus on statistically significant and high-confidence chromosomal interactions for downstream analysis, interactions called by *FitHiC2* were filtered to retain those with false discovery rates (FDR) below 0.05 for further analysis and 10^−5^ for visualization at genomic regions of interest. Significant interactions were visualized using the Integrated Genome Viewer (IGV) software (*134*). Loop sizes were not normally distributed and so compared using a Mann-Whitney-Wilcoxon two-sided test. Differential loops were detected by considering significant interactions that were present in only one of the conditions compared. To compare changes in the proportion of loops gained or lost between conditions, we used the Fisher’s exact test with false discovery rate (FDR) multiple test correction.

#### Functional categorization of chromatin loops

To study the biological roles of the chromatin loops, we compared their anchors to features defined from histone modifications, including “bivalent promoter”, “active promoter”, “active enhancer”, or “other” (regions lacking defined features), using *bedtools*. This approach enabled detailed categorization of chromatin loops based on genomic features intersecting with each loop anchor. Similarly, we categorised wild-type chromatin loops by the number of anchors (0, 1, 2) intersecting with MLL2 peaks (*35*), enabling the identification of MLL2-dependent loops for further inspection.

#### Gene ontology enrichment analysis of bivalent genes losing promoter-enhancer loops

Gene ontology (GO) enrichment analysis was performed using *g:Profiler* R client for genes that overlapped with “bivalent promoter” feature at their TSS and lost Bivalent Promoter-Active Enhancer chromatin loops in KO vs WT contrast at 72 hours. All genes intersecting with either WT or KO loops at 72 hours were used as the custom background, and significant GO terms were defined as entries with FDR < 0.05.

#### Aggregate peak analysis

Aggregate peak analysis (APA) was performed to visualise and compare contact signals for loops in each genomic feature or MLL2-binding category. Cool matrices were iteratively corrected using intrachromosomal contacts with *cooler balance* (*135*). Then, the expected Hi-C signals for each chromosome were calculated using *cooltools expected-cis* command (*136*). For each sample and category, pile-ups of observed over expected contact signals at the anchors of loop category were generated with *coolpup.py* using the balanced matrices (*137*). For consistency, wild-type loops were used as regions for visualization for all cell types. Heatmaps of aggregated signals were plotted using *plotpup.py* with additional parameters *--norm_corners* 10 to minimize differences in background signals. The comparison of signals across samples or categories was performed using *dividepups.py* command.

#### Integrating chromatin loop anchors with public protein binding profiles

To identify proteins enriched at loop anchor regions, we used the non-redundant dataset from Remap2022 public database (*65*) to obtain protein binding regions in mouse ESCs. To identify chromatin binding proteins enriched in one condition over another, we implemented a rank-based algorithm as follows: Statistically significant chromatin loops were categorised into discrete categories based on their presence across selected cell states using set operations. For all loops in each condition and for each chromatin binding protein, we counted the total number of binding events with significant loops at either end. Then, for each loop category, the proteins were ordered according to binding counts, and the relative enrichment score of each protein was calculated as the difference between its rank and the average rank across all other categories.

To explore the relationship between protein binding and their expression levels, we applied this algorithm for selected pairs of cells to generate a non-redundant list of top enriched proteins in cell-specific loops. The relative enrichment scores of these proteins were displayed against the log fold changes in expression of the corresponding genes using same contrast. For visualization purposes, we assigned negative values to the enrichment scores of proteins enriched in cell-specific loops from the baseline cell used in differential analysis.

#### Live-cell Single Molecule Localisation Microscopy data analysis

Data analysis of single-molecule localisations was carried out as previously described (*71*) with minor changes. Briefly, single-molecule localisations were extracted from 3D DHPSF movies using easyDHPSF software (*124*) in MATLAB (version 2016b) and the above-mentioned calibration file. After manual inspection of different thresholds, a relative localisation threshold of 80 and 120 was set for all 6 angles of H2B and *Sox2* datasets respectively. Individual localisations were assembled into trajectories using ‘trajectory-analysis’ (https://github.com/TheLaueLab/trajectory-analysis). Molecules in subsequent frames within a 500 nm radius were considered part of the same trajectory and hence connected, with a frame gap tolerance of 2 frames. To calculate the precision of the SMLM data, we conducted displacement analysis of fixed cells using the SMLM Analysis Toolbox (https://github.com/stanleystrawbridge/singleMoleculeLocalizationMicroscopyAnalysisToolBo <u>x</u>). Since displacements were not normally distributed, we compared them using a Kruskal-Wallis test and a post-hoc Dunn’s test with Holm correction.

A custom algorithm developed in our lab called ‘analyze-tracking-data’ was then deployed, which uses a sliding window along each trajectory to output four biophysical parameters for each sub-trajectory – anomalous exponent (α), diffusion coefficient (D_eff_), length of confinement (L_c_) and drift magnitude (*71*). Analysis was carried out using the following variable settings in the code: a sliding window of 7 frames for H2B and 11 frames for *Sox2*-loci, 0.5 s exposure time, a minimum trajectory length of 7 frames. Since chromatin mobility can be classified into two subpopulations, slow- or fast-moving chromatin, we also set this algorithm to use a 4-parameter Gaussian mixture model, considering all four biophysical parameters (default), to classify sub-trajectories as either slow- or fast-moving and to then output biophysical parameters separately for these two types of chromatin mobility. The mean parameter value for each replicate were plotted using custom scripts in RStudio (version 4.3.0). Since the mean parameter values were normally distributed, they were compared between conditions using an unpaired two-sided t-test.

## Author contributions

**Conceptualization:** SB and AFS. **Data curation**: LP, MS, AFS and SB. **Formal Analysis:** (scRNAseq) OD, JM, DA, SB, MB and BG. (Micro-C) LP, LM, SB and SAS. (Single-molecule localisation microscopy, SMLM) SY, MS and SB. (Quantitative immunofluorescence) MS. **Funding acquisition:** SB, MS and AFS. **Investigation:** (Neuroectoderm differentiation, ChIP-qPCR) KN, AK, KA and AFS. (scRNAseq) JM and MS. (Micro-C) MS with help from AW and GA. (SMLM) MS and GA. **Methodology:** No new methods used. **Project administration:** SB, BH and AFS. **Resources:** (*Mll2* knockout, *Mll2-GFP* and *Mll2* NCat cell lines) KN, AK, JF, KA and AFS. (Cell lines expressing HaloTag-H2B and dCas9-GFP) MS and GA. (SMLM instrument alignment and maintenance) ZZ and DK. **Software:** (SMLM) SES, AW, SY and DH. **Supervision:** SB, BH and AFS. **Validation:** (Quantitative immunofluorescence) MS, OD and NC. **Visualization:** (scRNAseq and Micro-C) DA, OD, JM and LP. (Immunofluorescence) KN, AK and MS. (SMLM) MS and SY. **Writing – original draft:** SB and AFS. **Writing – review & editing:** MS, OD and AW.

## Acknowledgements

We thank all the members of the Basu and Stewart labs, and Michelle Meredyth, for insightful and critical discussions. We thank the Genomics Core Facility, Cancer Research UK Cambridge Research Institute for 10x Chromium single cell library preparation and Illumina sequencing. We thank the Cambridge Stem Cell Institute (CSCI) Genomics Core Facility for assisting with quality control and submission of sequencing libraries. We thank Peter Humphreys, Darran Clements and Louis Elfari in the CSCI Imaging Facility for training students and assisting with experiments. We thank Sally Lees and the CSCI Tissue Culture Core Facility for technical advice and support. We thank Arash Shahsavari, Elze Lauzikaite and Irina Mohorianu in the CSCI Bioinformatics Core Facility for help pre-processing of some of the sequencing datasets. The PBase and dCas9-GFP plasmids were kind gifts from Dr Brian Hendrich and Prof Joanna Wysocka. We also thank Menan Loganathan and Igor Orsine de Almeida for preliminary analyses.

MS and AW were funded by the Wellcome Trust PhD programme in Stem Cell Biology and Medicine (224929/Z/22/Z and 218481/Z/19/Z). OD was funded by the Addenbrooke’s Charitable Trust and the UKRI Medical Research Council (MR/Y000463/1). LP and GA were funded by the UKRI Biotechnology and Biological Sciences Research Council (BB/W000423/1, BB/W000423/2). AK, KA and AFS were funded by Deutsche Forschungsgemeinschaft grants (STE 903/12-2 and STE 903/13-1). DA was funded by the Simons Foundation Autism Research Initiative. JM and SB were funded at the Cambridge Stem Cell Institute through a starting grant from the Wellcome Trust (203151/Z/16/Z) and UKRI Medical Research Council (MC_PC_17230). SB was also funded by the Trinity College Stem-Cell Medicine Senior Postdoctoral Researcher Fund and Imperial College London. SES was funded by the Joint Research Grant (Isaac Newton Trust, Wellcome Trust ISSF, University of Cambridge), the CSCI Wellcome Trust seed fund and the Wellcome Trust Sir Henry Wellcome Fellowship (224070/Z/21/Z).

## Materials availability

All plasmids and cell lines generated in this study are available from Addgene or from the lead contact with a completed materials transfer agreement.

## Data and code availability

All imaging datasets are available from the lead contact. The XYZT single molecule trajectory data files are uploaded to Zenodo and will be made available upon publication. The scRNAseq and Micro-C datasets reported in this study are available from the Gene Expression Omnibus (GEO) repository under accession codes GSE276814 and GSE279709. All code used is already open-source and links have been provided to the relevant websites. Original code developed for this study, the SMLM Analysis Toolbox, has been made available on Github: (https://github.com/stanleystrawbridge/singleMoleculeLocalizationMicroscopyAnalysisToolBox).

---

**Fig. S1.**
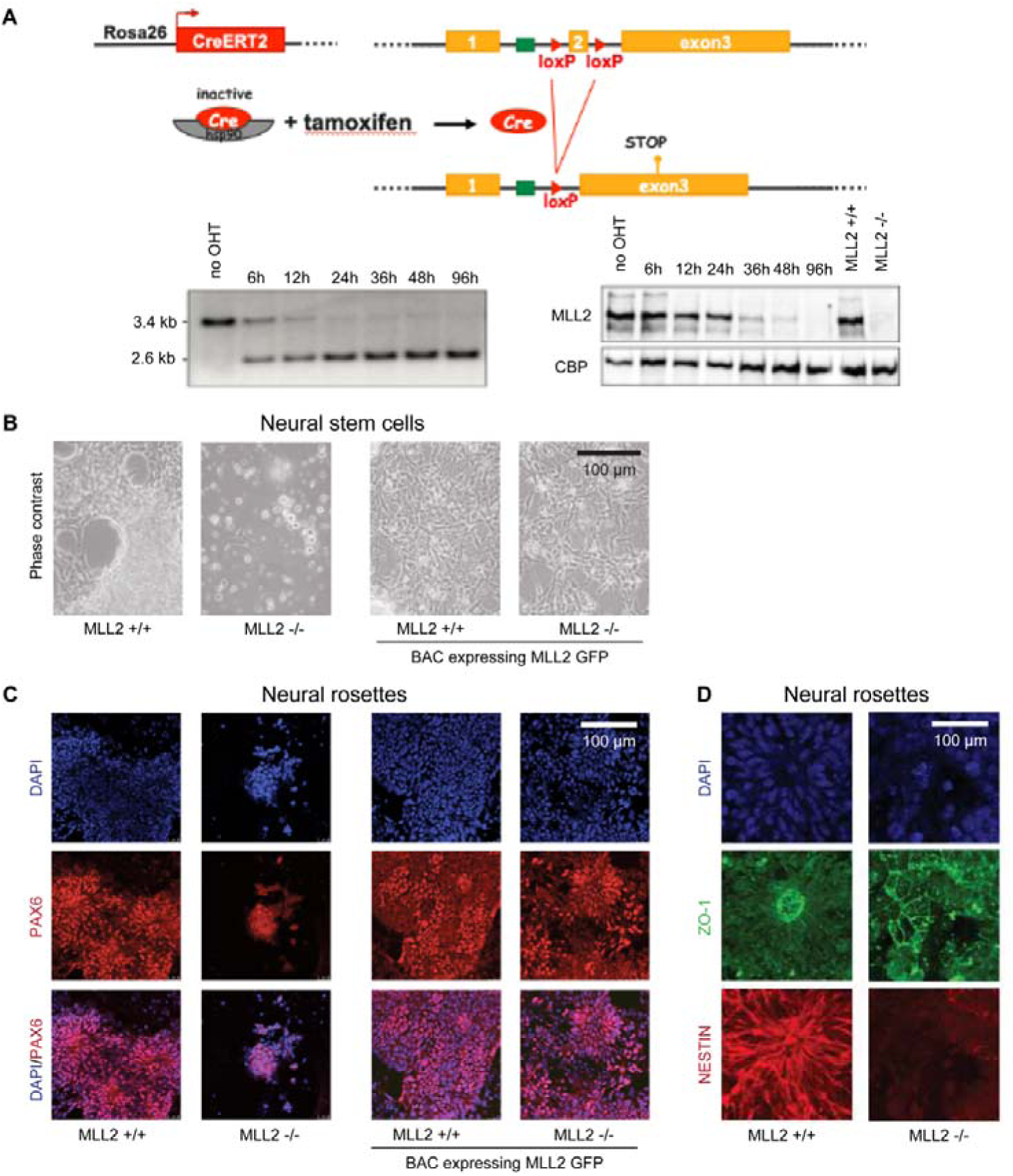
MLL2 knockout disrupts neuroectoderm specification. **(A)** (Top) Schematic representation of the conditional knockout (cKO) strategy. *Mll2* cKO cell line has loxP sites either side of *Mll2* exon 2 and expresses CreERT2 from a Rosa26 promoter to allow tamoxifen-induced removal of *Mll2* exon 2. (Bottom left) Genomic PCR confirms removal of *Mll2* exon 2 within 24 hr. (Bottom right) Western blot confirms depletion of MLL2 protein within 48 hr. **(B)** Phase contrast images at the neural stem cell stage of control (MLL2 +/+) and *Mll2* cKO (MLL2−/−) cells show that *Mll2* cKO cells fail to generate neural stem cells. **(C)** DAPI stain (blue), PAX6 expression (red) and merge at the neural rosette stage in control (MLL2 +/+) and tamoxifen-induced *Mll2* cKO cells (MLL2 −/−) but also control and *Mll2* cKO cells expressing exogenous GFP-tagged MLL2 from an integrated bacterial artificial chromosome (BAC). **(D)** DAPI stain (blue), ZO-1 expression (red) and NESTIN expression (red) at the neural rosette stage in control (MLL2 +/+), tamoxifen-induced MLL2-deficient (MLL2 −/− cells).

**Fig. S2.**
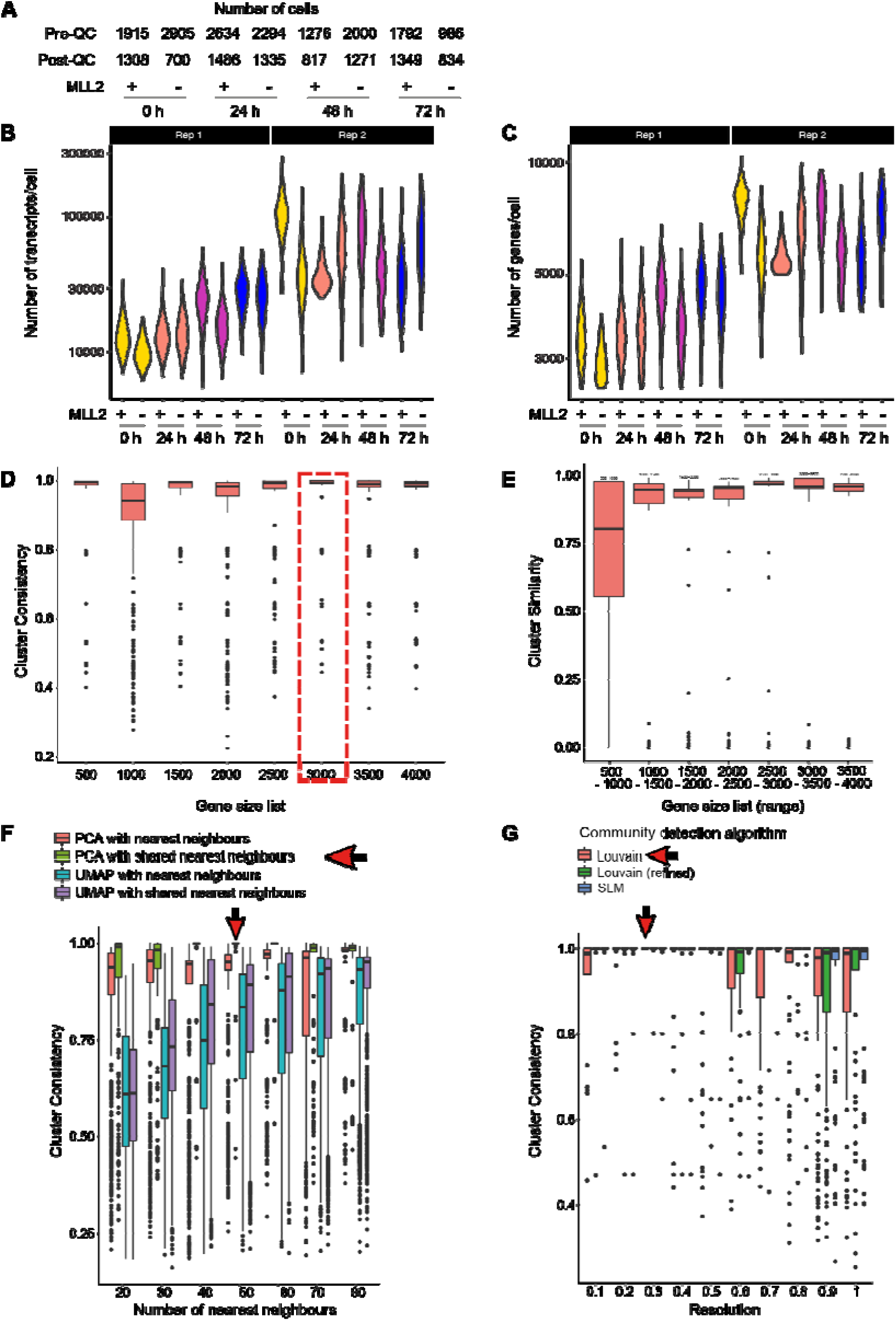
Quality control of 10x scRNAseq datasets and cluster analysis. **(A)** Table containing number of cells per sample. **(B)** Violin plot showing number of transcripts/cell post filtering [see **(A)** for sample size]. **(C)** Violin plot of number of genes per cell from 10x scRNAseq post-filtering [see **(A)** for sample size]. **(D-E)** The effect of gene list size (of the most variable genes), used in the PCA dimensionality reduction step, on **(D)** cluster consistency and **(E)** similarity (n = 100 iterations of randomly sampled genes). **(F)** The effect of different base embeddings (PCA vs UMAP), nearest neighbouring algorithm (standard or shared) and number of neighbours on clustering consistency. **(G)** The effect of different community detection algorithms on cluster consistency. **(D-G)** red box and arrow indicate the parameters selected for scRNAseq analysis. (n = 100 iterations of randomly sampled genes)

**Fig. S3.**
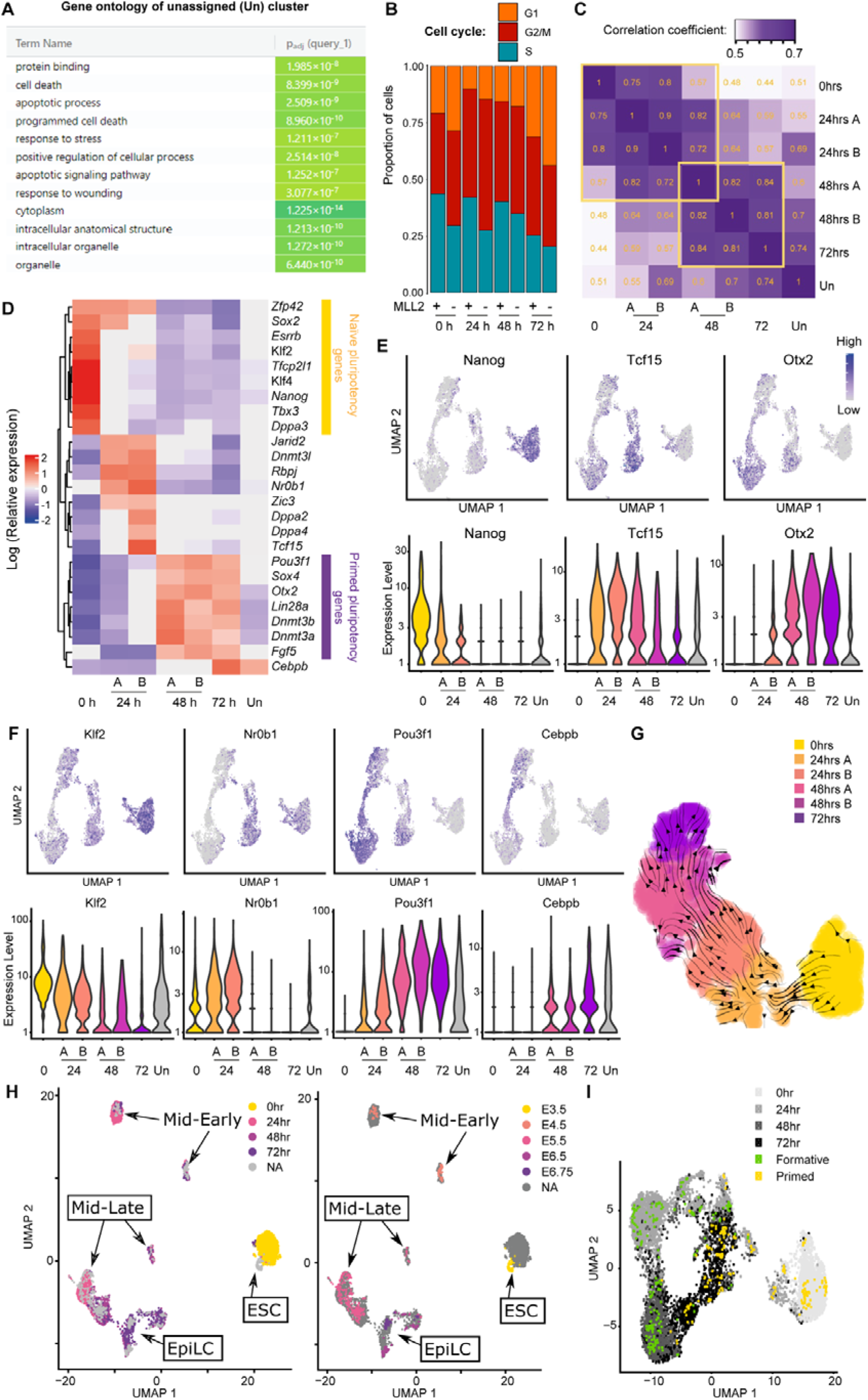
Cluster assignment and cluster-specific genes. (**A**) G profiler output for unassigned cluster shows a signature of apoptosis and programmed cell death. **(B)** Cell cycle analysis show significant changes between wild-type and *Mll2* conditional knockout (cKO) cells with more cells in G1 and fewer cells in S phase [see **(Fig. S2A)** for sample size]. **(C)** Spearman correlation of all genes assigned to the second highest expression quartile instead of only variable genes used for dimensionality reduction. **(D)** Heatmap of annotated gene set showing relative gene expression in assigned clusters. The gene set is based on previous publications of genes involved in pluripotent cell priming. **(E-F)** Single-cell expression of naïve, intermediate and primed genes from **Figure 2B** +/− *Mll2* for **(E)** well-known genes and **(F)** less well-known genes. Expression represented as (top) a UMAP plot coloured by transcript level (grey to blue for low to high expression) and (bottom) as violin plots of transcript level per assigned cluster. Number of cells: 1215/638 (0 hr +/− MLL2), 428/532 (24 hr A +/− MLL2), 998/671 (24 hr B +/− MLL2), 759/914 (48 hr A +/− MLL2), 133/223 (48 hr B +/− MLL2), 348/343 (72 hr +/− MLL2) and 716/819 (Unassigned). **(G)** UMAP plot regenerated from **Figure 2B** after removal of unassigned cluster and using parameters that better represent distance between cells rather than between clusters. Cells in UMAP arranged by pseudotime metric calculated from CellRank pipeline and with arrows indicating directionality. **(H)** UMAP plot combining our data (n = 8737 cells) with published mouse embryo *in vivo* datasets (n = 721 cells) (*48*). **(I)** UMAP plot combining our data (n = 8737 cells) with published *in vitro* datasets of formative-like (n = 192 cells) and primed ESCs (n = 192 cells) (*40*).

**Figure S4.**
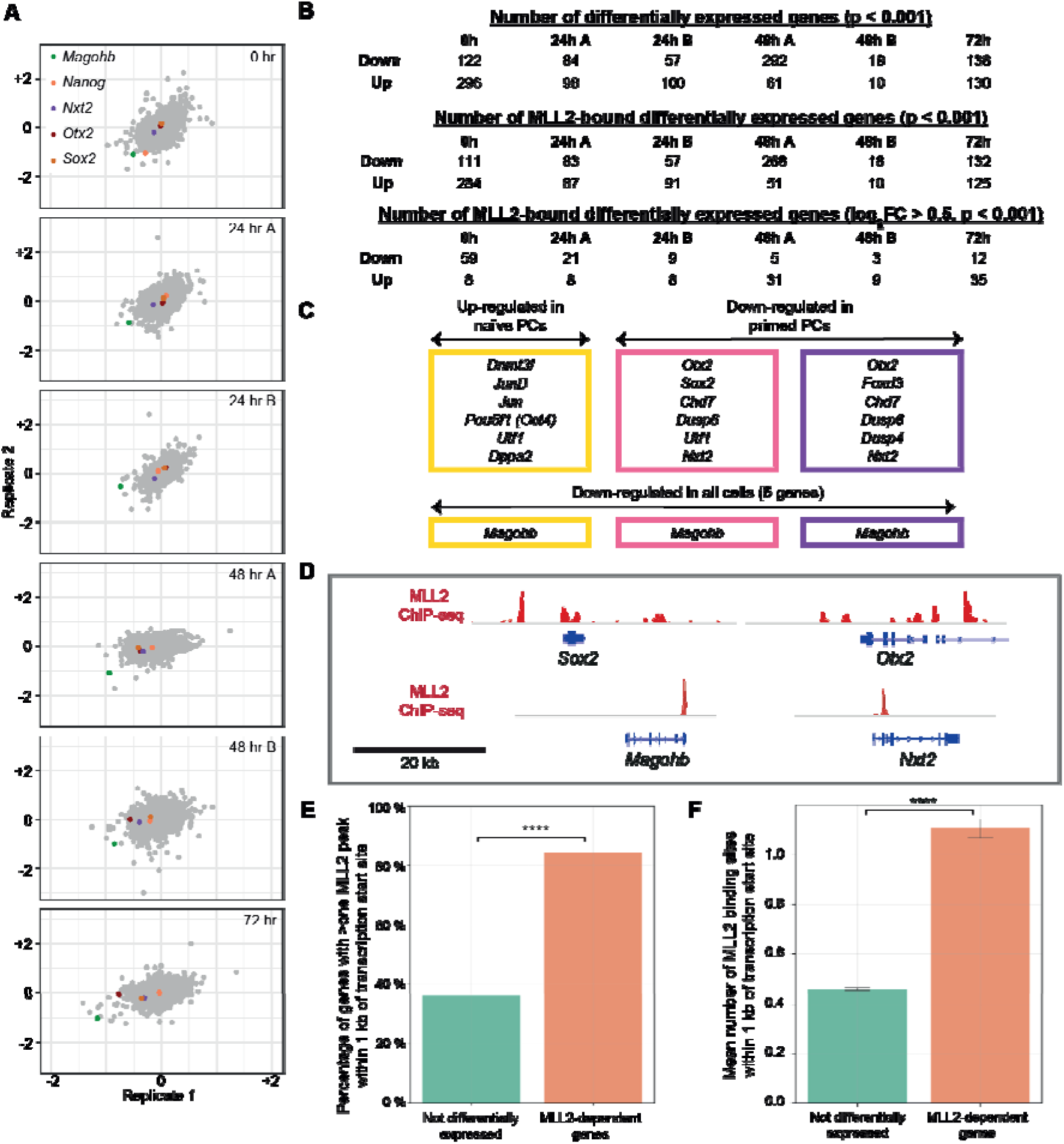
Identification of MLL2-dependent genes. **(A)** Comparison of scRNAseq replicates show reproducibility in log_2_FC for *Mll2*-dependent genes. **(B)** Table showing number of DEGs (top) and number of DEGs with MLL2 bound within 1 kb of the gene’s transcription start site: log_2_FC > 0 (middle) or log_2_FC > 0.5 (bottom). **(C)** Table showing example DEGs up-regulated in naïve PCs and down-regulated in primed PCs. **(D)** UCSC genome browser tracks showing MLL2 ChIP-seq (*35*) for the *Mll2*-dependent genes *Sox2*, *Otx2*, *Magohb* and *Nxt2*. **(E)** Bar chart showing an increase in the proportion of genes bound by MLL2 at differentially expressed genes versus at those with no transcriptional change. [Chi-squared test: p = 10^−267^] **(F)** Bar chart showing an increase in the mean number of MLL2 peaks at differentially expressed genes versus at those with no transcriptional change. [error bars represent 95 % confidence interval, Mann Whitney Test: p = 10^−273^]

**Figure S5.**
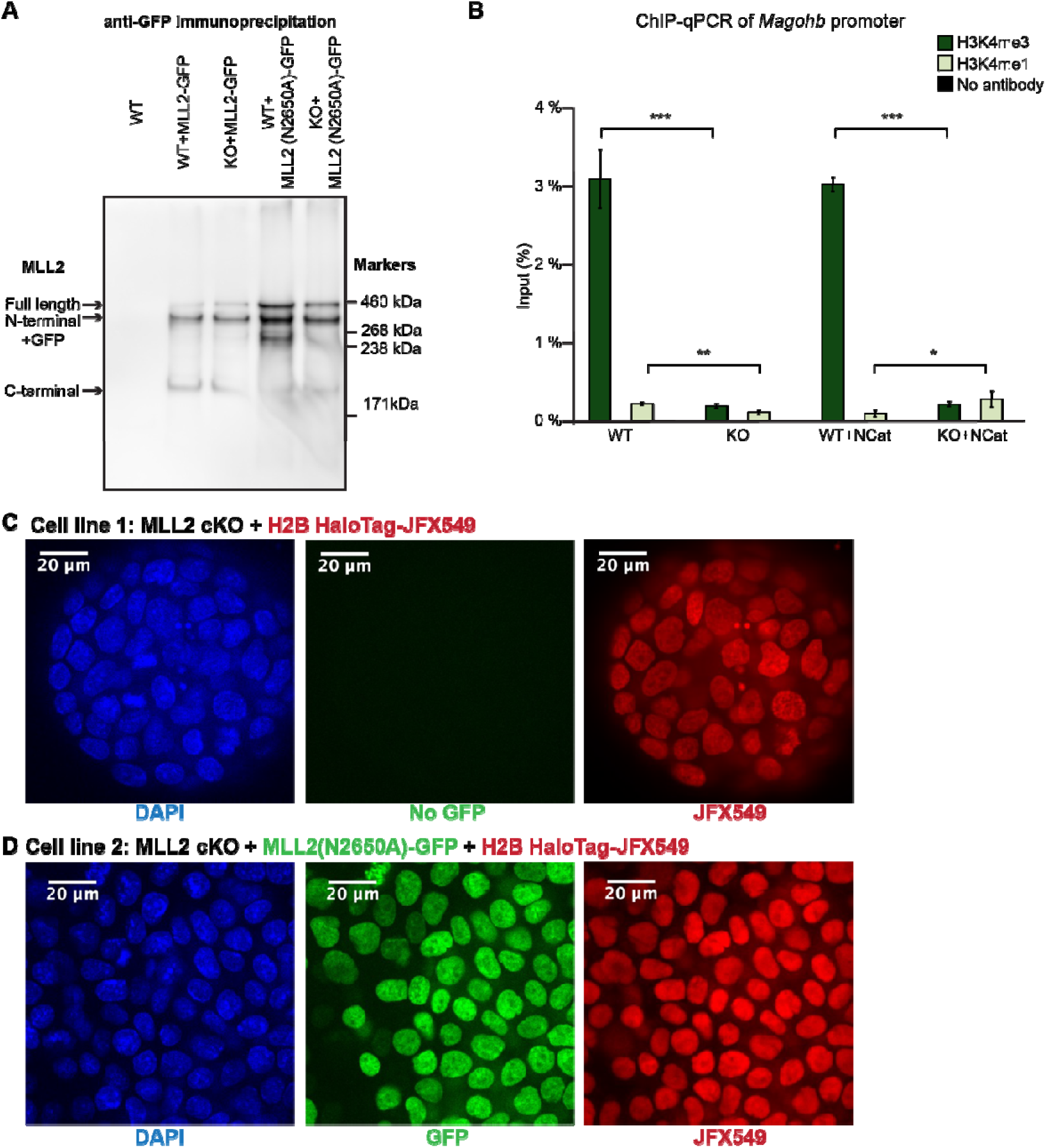
Generation of cell lines expressing catalytically-dead MLL2 and/or HaloTag-tagged histone H2B. **(A)** Immunoprecipitation using anti-GFP followed by SDS PAGE gel shows expression of GFP-tagged MLL2. **(B)** ChIP-qPCR at *Magohb* gene promoter for H3K4me3 and H3K4me1 [unpaired two-sided t-test, p-values: ***10^−4^ (H3K4me3 WT vs KO), **10^−3^ (H3K4me1 WT vs KO), ***10^−4^ (H3K4me3 WT + NCat vs KO + NCat), *0.04 (H3K4me1 WT + NCat vs KO + NCat)] **(C)** Representative images of *Mll2* conditional knockout (cKO) cell line expressing HaloTag-tagged histone H2B, where histone H2B molecules were labelled using JFX_549_ dye (red) and imaged alongside DAPI (blue) to confirm nuclear localisation. **(D)** Representative images of *Mll2* cKO cell line ectopically expressing a GFP-tagged catalytically-dead form of MLL2 and HaloTag-tagged histone H2B, where histone H2B molecules were labelled using JFX_549_ dye (red) and imaged alongside DAPI (blue) and GFP-tagged MLL2 (green) to confirm expression of catalytically-dead form of MLL2 and nuclear localisation.

**Figure S6.**
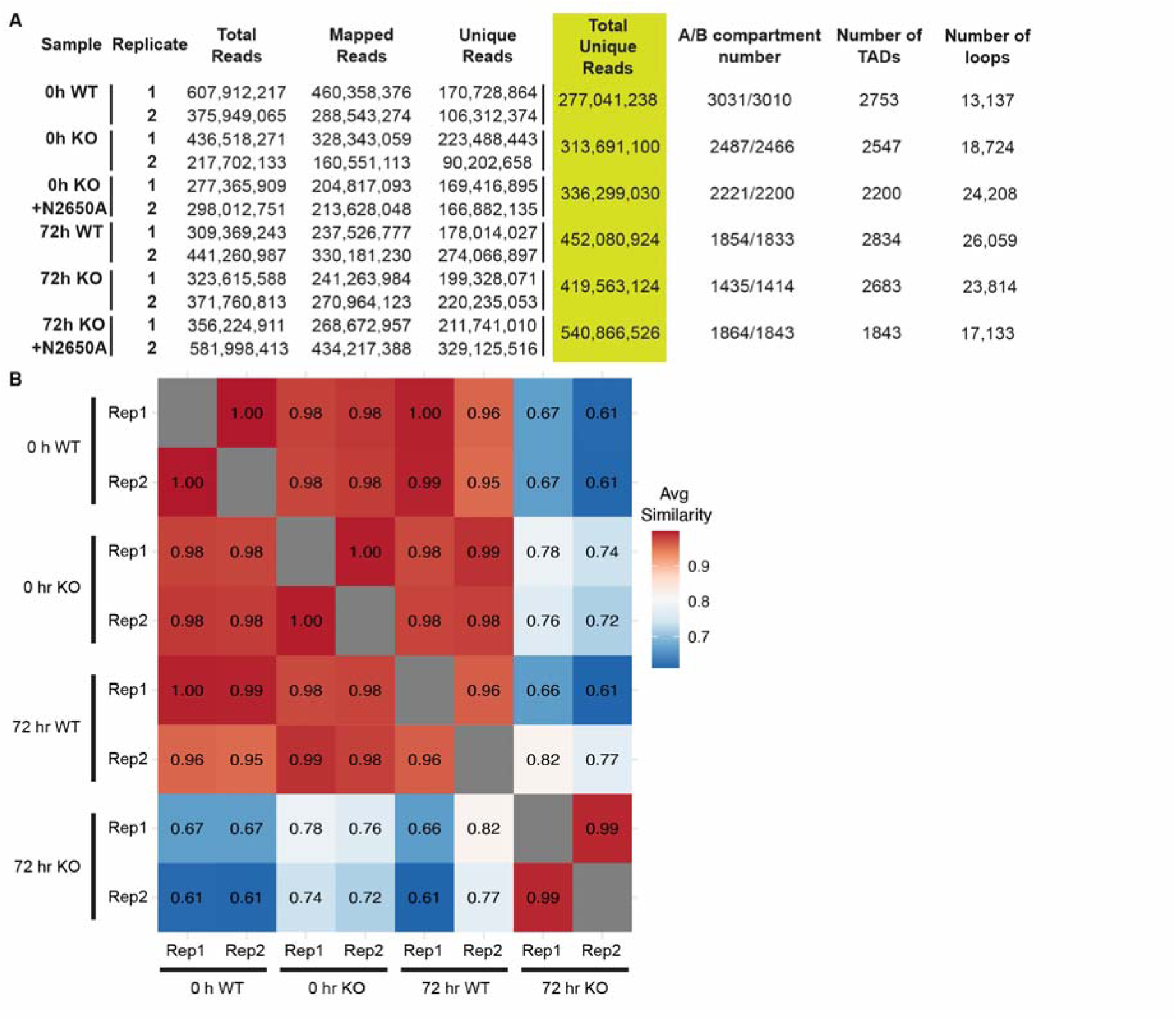
Quality control of Micro-C analysis. **(A)** Table showing number of reads, 3D looping contacts, compartments, TADs and significant loops detected by FitHiC2 (q<0.05). **(B)** HICRep shows consistency between replicates when comparing contacts at a range of bin sizes (0.25, 0.5, 1 and 2.5 Mb).

**Figure S7.**
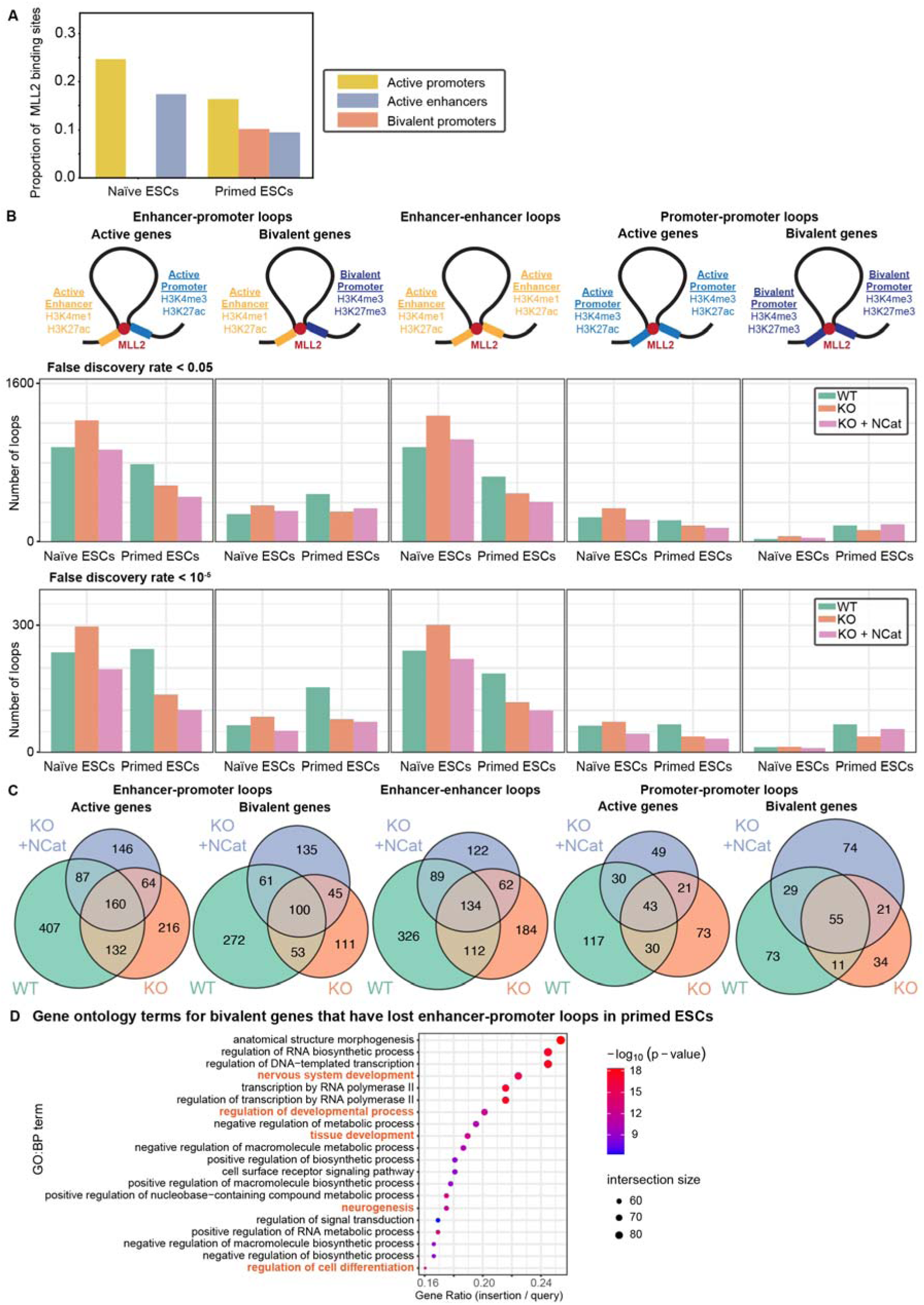
Enhancer-promoter analysis shows that MLL2 knockout decreases looping at enhancers and promoters which is rescued at bivalent genes by expression of MLL2 NCat. **(A)** Proportion of MLL2 binding sites that overlap with active enhancers, active promoters and bivalent promoters in wild-type (WT) naïve and primed ESCs. Enhancers are defined by overlapping H3K4me1 and H3K27ac peaks, active promoters by overlapping H3K4me3 and H3K27ac peaks and bivalent promoters by overlapping H3K4me3 and H3K27me3 peaks. Peaks identified from WT cells (*51*). **(B)** (Row 1) Schematic showing how categories of loops are defined as enhancer-promoter loops at active genes; enhancer-promoter loops at bivalent genes; enhancer-enhancer loops; promoter-promoter loops at active genes; or promoter-promoter loops at bivalent genes. (Row 2-3) Barplots showing number of loops calculated with a false discovery rate (FDR) of (Row2) 0.05 or (Row 3) 10^−5^ when comparing WT, *Mll2* cKO (KO) and KO expressing catalytically-dead MLL2 (KO+NCat) for both naïve and primed ESCs. **(C)** Euler diagram showing overlap between WT (green), KO (orange and KO+NCat (pink) conditions shows that loops recovered in KO+NCat conditions are not always the same as in WT cells. **(D)** GO analysis shows that bivalent genes with loop changes in primed ESCs are associated with nervous system development.

**Figure S8.**
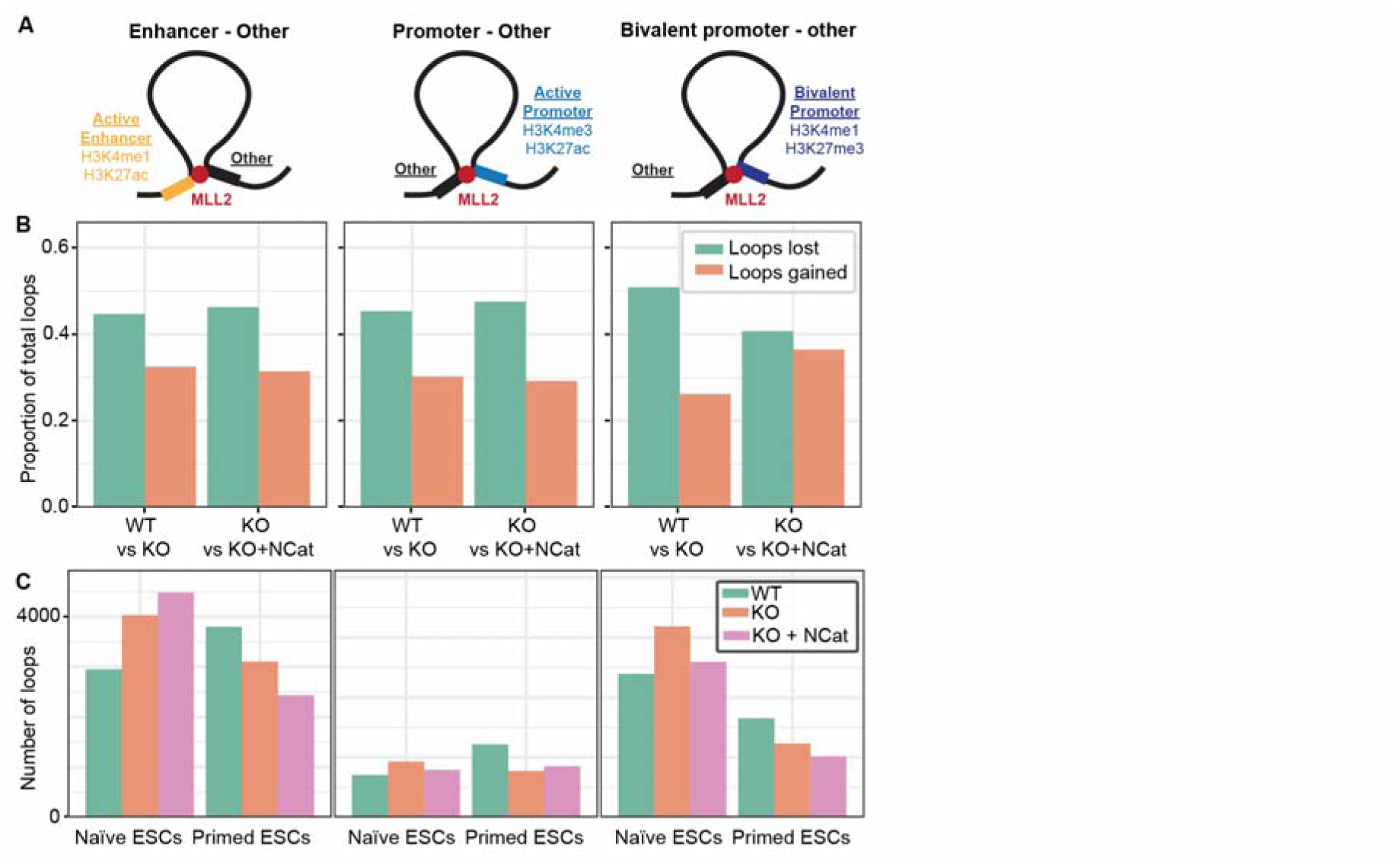
Enhancer-promoter analysis shows that MLL2 knockout increases non-specific looping at enhancers and promoters. **(A)** Schematic showing how categories of non-specific loops are defined as enhancer-other loops and promoter-other loops at active or bivalent genes. **(B)** Barplot showing proportion of non-specific loops gained (green) and lost (orange) in primed ESCs when comparing wild-type (WT) to *Mll2* conditional knockout (KO) or KO to KO expressing catalytically-dead MLL2 (KO+NCat) [Fisher’s exact test with false discovery rate (FDR) multiple test correction, p-values: 10^−40^/10^−47^, 10^−39^/10^−43^ and 10^−126^/10^−4^ (E-other, P-other and other-other loops in WT vs KO/ KO vs KO+NCat)]. **(C)** Barplot showing number of non-specific loops when comparing WT (green), KO (orange) and KO+NCat (pink) conditions.

**Figure S9.**
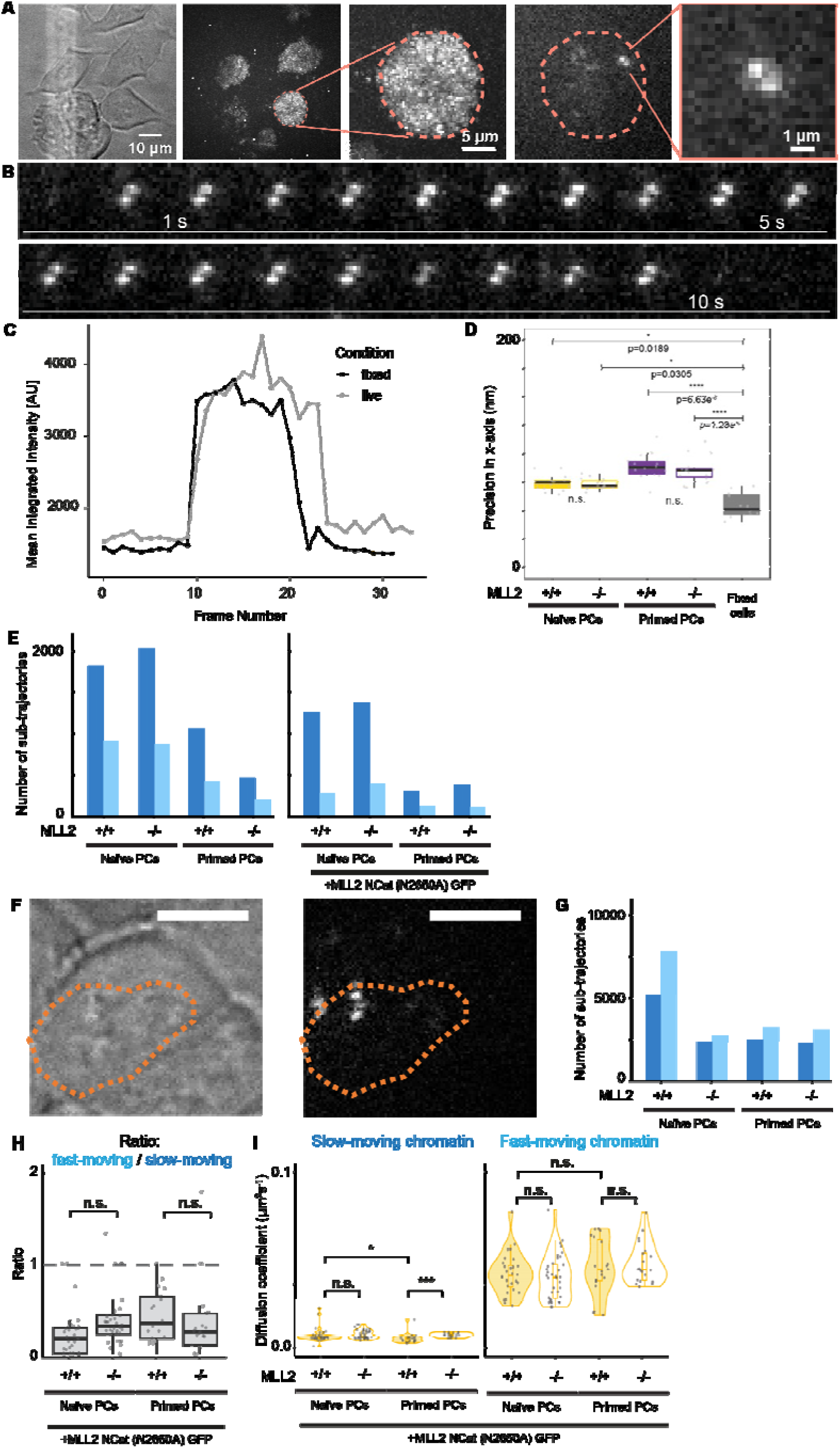
Control experiments for SMLM. **(A)** (Left to right) Bright-field image of a field of view containing 3-4 cells. Next, we have a sum project of a single-molecule localisation microscopy (SMLM) video showing nuclear localisation of histone H2B within 3-4 nuclei. We then show a zoom of one of these nuclei with the nuclear periphery indicated by an orange dotted line. Then we show a representative SMLM image with a single PA-JF_549_-HaloTag-tagged histone H2B molecule. We then show a zoom of the single molecule in the final panel. **(B)** Representative time-lapse montage showing single-step appearance and disappearance of PA-JF_549_-HaloTag-tagged histone H2B molecule over 11 s (or 22 frames) taken from a live-cell SMLM video. **(C)** Intensity trace of histone H2B molecules labelled using PA-JF_549_ dye and tracked at 500 ms time resolution show single-step photobleaching. **(D)** Distance travelled by H2B between frames significantly higher in live versus fixed cells along the x, y and z directions [Kruskal-Wallis test and a post-hoc Dunn’s test with Holm correction, p-values indicated on plots]. **(E)** Number of trajectories in H2B samples. **(F)** Representative bright-field and fluorescence image of cells labelled for imaging of *Sox2* gene using deactivated Cas9 and Atto647-labelled gRNAs. **(G)** Number of trajectories in *Sox2* gene tracking samples. **(H)** Proportion of H2B molecules observed in slow- and fast-moving chromatin states in WT/KO cells expressing N2650A catalytically-dead form of MLL2. Dots represent ratios calculated from independent videos (n>25 videos, 3-5 cells per video) [unpaired two-sided t-test, p-values: 0.14 (WT+NCat vs KO+NCat in naïve PCs), 1.0 (WT+NCat vs KO+NCat in primed PCs)] (**I**) Box-and-violin plot showing apparent diffusion coefficient D_app_ of (left) slow- and (right) fast-moving chromatin for naïve and primed PCs in WT/KO cells expressing N2650A catalytically-dead form of MLL2. [unpaired two-sided t-test, p-values: *0.02 (slow-moving WT+NCat naïve vs primed PCs), 0.2 (slow-moving WT+NCat vs KO+NCat in naïve PCs), ***10^−4^ (slow-moving WT+NCat vs KO+NCat in primed PCs), 1.0 (fast-moving WT+NCat naïve vs primed PCs), 1.0 (fast-moving WT+NCat vs KO+NCat in naïve PCs), 1.0 (fast-moving WT+NCat vs KO+NCat in primed PCs)]

**Figure S10.**
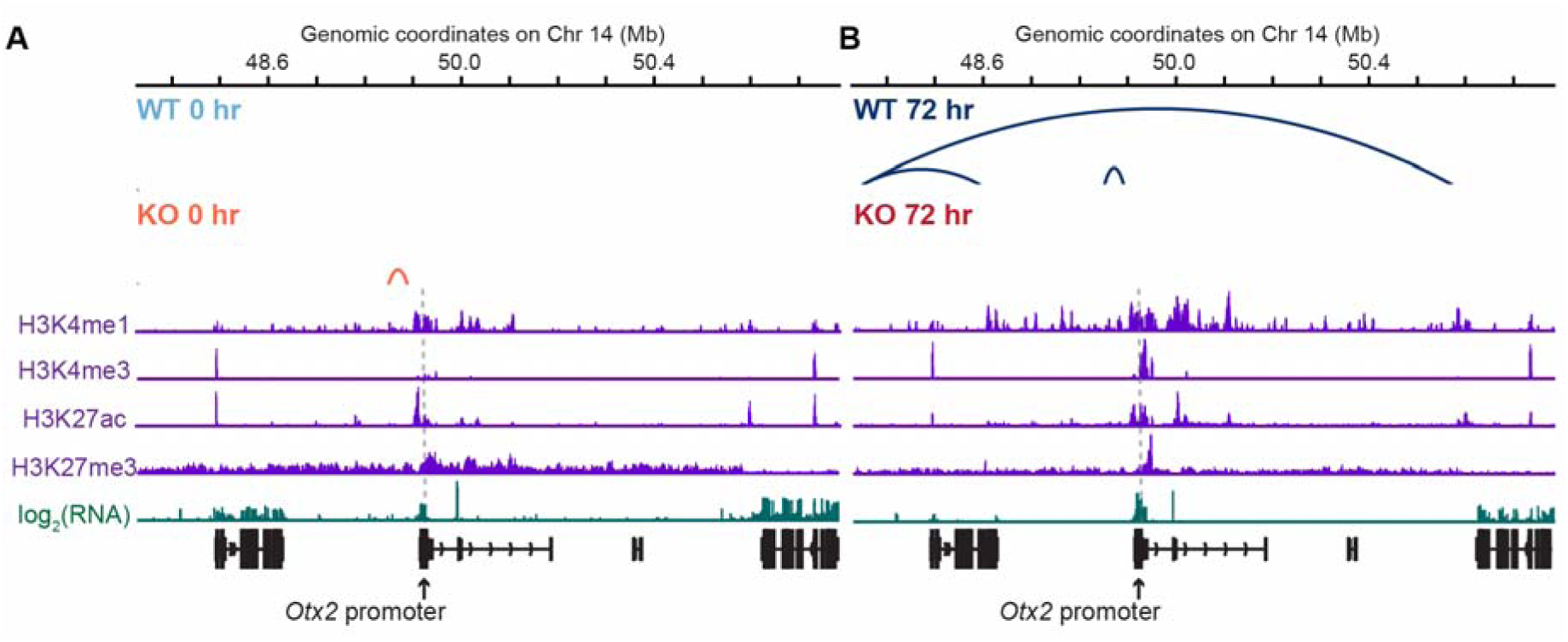
MLL2 knockout drives gene-specific and cell type-specific changes in 3D chromatin looping at *Otx2* gene. **(A-B)** High-confidence loops detected by FitHiC2 (FDR<10^−5^) alongside published RNA-seq and ChIP-seq of histone marks (*16*) at *Otx2* gene in (**A**) naïve and (**B**) primed ESCs for WT and KO cells. Grey dotted line represents *Otx2* promoter. Grey loops are common to both conditions, light blue to WT naïve ESCs, light red to KO naïve ESCs, dark blue to WT primed ESCs and dark red to KO primed ESCs (FDR<10^−5^).

